# High-Dimensional Gene Expression and Morphology Profiles of Cells across 28,000 Genetic and Chemical Perturbations

**DOI:** 10.1101/2021.09.08.459417

**Authors:** Marzieh Haghighi, Juan Caicedo, Beth A. Cimini, Anne E. Carpenter, Shantanu Singh

**Affiliations:** Broad Institute of MIT and Harvard

## Abstract

Cells can be perturbed by various chemical and genetic treatments and the impact on the cells’ gene expression (transcription, i.e. mRNA levels) and morphology (in an image-based assay) can be measured. The patterns observed in this high-dimensional profile data can power a dozen applications in drug discovery and basic biology research, but both types of profiles are rarely available for large-scale experiments. Here, we provide a collection of four datasets with both gene expression and morphological profile data useful for developing and testing multi-modal methodologies. Roughly a thousand features are measured for each of the two data types, across more than 28,000 thousand chemical and genetic perturbations. We define biological problems that use the shared and complementary information in these two data modalities, provide baseline analysis and evaluation metrics for multi-omic applications, and make the data resource publicly available (http://broad.io/rosetta).

## Introduction

**B**iological systems can be quantified in many different ways. For example, researchers can measure the morphology of a cell using microscopy and image analysis, or molecular details such as the levels of messenger RNA (mRNA) or protein in cells. Historically, biologists chose a single feature to measure for their cell samples, based on their prior knowledge or hypotheses. Now, “profiling” experiments capture a high-dimensional profile of features for each sample, and hundreds to thousands of samples can be quantified. This allows the discovery of unexpected behaviors of the cell system.

Profiling experiments carried out at large scale remain expensive, even for a single profiling modality. We observed that no public dataset exists providing both genetic and chemical perturbation of cells with two different kinds of profiling readouts. Such a dataset would enable multi-modal (also known as *multi-omic*) analyses and applications. Examples include integrating the two data sources to better predict a compound’s activity in an assay ^1^, predicting the mechanism of action of a drug based on its profile similarity to well-understood drugs ^2^, or predicting a gene’s function based on its profile similarity to well-understood genes ^3^.

Observing a system from multiple perspectives is known to reveal patterns in data that may not be visible in individual perspectives. Machine learning methods have been explored in various fields to learn from multiple sources to make better inferences from data ^4^. In biology, the advancement of technologies for measuring multi-omics data has sparked research investigating the relationship and integration of different high-dimensional readouts ^5^. For example, transcriptomics, proteomics, epigenomics and metabolomics data can be combined to predict the mechanisms of action (MoAs) of chemical compounds ^6^.

Here, we created a collection of gene expression and morphology datasets with the scale and annotations needed for machine learning research in multi-modal data analysis and integration. This Resource provides two different, rich views on the cells by providing roughly a thousand mRNA levels and a thousand morphological features when samples of cells are perturbed by hundreds to thousands of different conditions, including chemical and genetic. Furthermore, we present a framework for thinking about the utility of multi-modal data by defining applications where the shared information, and the complementary information, across data types can be useful, using terminology understandable to those new to the biological domain. We demonstrate example applications within each group, uncover interesting biological relationships, and provide baseline methods, code, evaluation metrics, and benchmark results for each, as a foundation for future biologically-oriented machine learning research.

## Results

### Gene expression and morphological profiles

All datasets were created at our institution (see Supplementary 1) and involved one of two types of “inputs”: chemical perturbations and genetic perturbations (Figure 1). There are also two types of high-dimensional outputs measured: gene expression profiles and morphological profiles, each with roughly 1000 features measured. For each of the datasets, in a single laboratory, cells are plated into two sets of identical plates, each plate gets treated with chemical (or genetic) perturbations identically, and then one set is used to measure gene expression and the other set to measure morphology.

**Figure 1.**
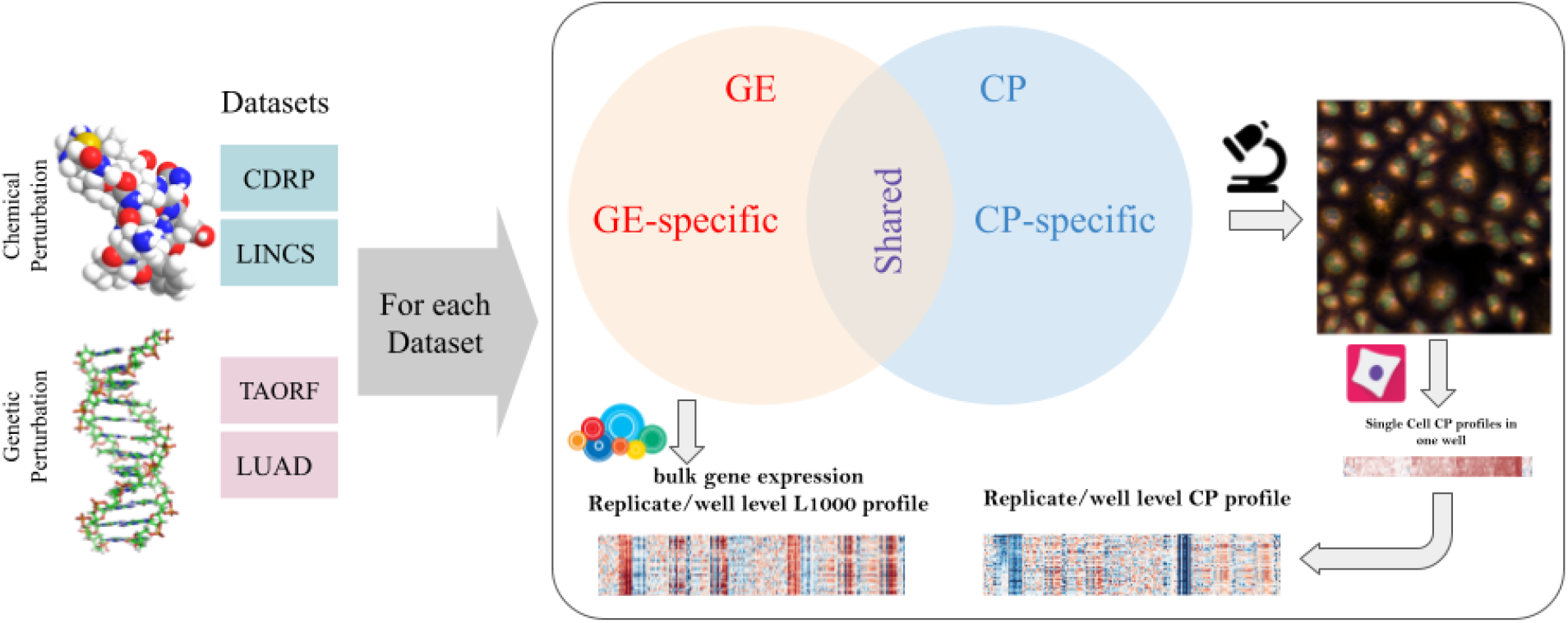
Multi-modal datasets overview. Multi-modal genetic and chemical perturbation datasets are valuable for many applications. For each dataset, Cell Painting (CP) and Gene Expression (GE) L1000 assays were used to collect morphological and gene expression representations (profiles), respectively. The datasets are described in Supplementary 1. Credits: Chemical structure by GJ Owns, distributed under a CC0 3.0 license. DNA structure by Richard Wheeler, distributed under a CC-BY-SA 3.0.

We captured gene expression (GE) (mRNA profiles) using the L1000 assay ^7^. The levels of mRNA in the cell are often biologically meaningful - collectively, mRNA levels for a cell are known as its transcriptional state. The L1000 assay reports a sample’s mRNA levels for ~978 genes at high-throughput, from the bulk population of cells treated with a given perturbation. These “landmark” genes capture approximately 82% of the transcriptional variance for the entire genome^7^; the specific genes’ mRNAs that are measured can be different across datasets, though largely overlapping (see Methods). We note that “genes” are an input (individual genes are overexpressed as the perturbation in some datasets) and an output (gene expression profiles are the measured mRNA levels for each landmark gene in the L1000 assay); this can cause confusion for researchers new to the domain.

We captured morphological profiles using Cell Painting (CP) ^8^. This microscopy assay captures fluorescence images of cells colored by six well-characterized fluorescent dyes to stain the actin cytoskeleton, Golgi apparatus, plasma membrane, nucleus, endoplasmic reticulum, mitochondria, nucleoli, and cytoplasmic RNA in five channels of high-resolution microscopy images. Images are processed using CellProfiler software ^9^ to extract thousands of features of each cell’s morphology such as shape, intensity and texture statistics, thus forming a high-dimensional profile for each single cell. The profiles are then aggregated (population-averaged) for all imaged single cells in each sample well.

For both data types, aggregation of all the replicate-level (equivalent to well-level) profiles of a perturbation is called a treatment-level profile. In our study, we used treatment-level profiles for each perturbation in all experiments but have provided replicate-level profiles for researchers interested in further data exploration. Therefore, in our experiments, each perturbation in each dataset has two corresponding vectors of measurements for each modality; one treatment-level profile for gene expression and one treatment-level profile for morphological measurements. Gene expression and morphological measurements are taken from two different sets of plates; there is therefore no direct, one-to-one correspondence between the two readouts at the replicate level. We also note that of the eight datasets provided (four datasets x two modalities), four have been the subject of previous research published by researchers at our Institute ^3,10,11^; here we complete the matrix by providing the missing data type for each pair, organizing them, and providing benchmarks.

### Shared versus complementary information content

Cell morphology and gene expression are two very different kinds of measurements about a cell’s state, and their relationship is known to be complex. For example, a change in morphology can induce gene expression changes ^12^ and gene expression changes can induce a change in cell morphology ^13,14^. However, a strict relationship is not always the case; many drugs impact cells’ mRNA *or* morphology profile, but not both ^10,15,16^. For example, changes in protein stability or post-translational modifications can induce changes in morphology without changes in gene expression; the Rho-family small GTPases are one example where morphology changes on a timescale much too short to be explained by changes in mRNA ^17^. Furthermore, the two data types are collected at different timepoints, determined as optimal for each individually. Therefore, even if technical artifacts were non-existent, we do not expect a one-to-one map between these two modalities. We therefore hypothesize that the information in each data type consists of a shared subspace, a modality-specific complementary subspace, and noise (Figure 1). Both subspaces can be exploited for biological applications.

### Shared subspace across two modalities

The shared subspace between gene expression and cell morphology is beginning to be explored. For example, cross-modal autoencoders learned the shared latent space for single-cell RNA-seq and chromatin images in order to integrate and translate across modalities ^18^. In another study, probabilistic canonical correlation analysis learned a shared structure in paired samples of histology images and bulk gene expression RNA-seq data, suggesting that shared latent variables form a composite phenotype between morphology and gene expression that can be useful ^19^. Many uncovered relationships will not be transferable from one experimental batch to another, particularly if great differences exist: for example, histology images differ in many ways from fluorescence microscopy images, yet some features, such as nuclear shape, might be consistent across different experimental techniques.

The existence of a shared subspace enables multiple applications. Most prominently, if sufficient shared information is present, one modality can be computationally predicted (i.e. inferred, estimated) using another, saving significant experimental resources. For example, one could predict the expression level of genes of interest given their morphological profiles from already-available images, even from patients whose samples are no longer available for mRNA testing. Or, one could generate images or morphological profiles from large libraries of mRNA profiles.

Another use of shared subspace is to identify relationships between specific features of the two types. For example, a morphological feature and a specific gene’s mRNA level may be tightly linked, which can yield clues as to the biological mechanisms underlying their relationship. As well, inspecting *which* genes can be well-predicted may shine light on general relationships between mRNA levels and morphology for different classes of genes^20^; enrichment analysis of these groups of genes could also lead to biological pattern discoveries. Researchers have used linear regression and enrichment analysis to explore the association between variations in cell morphology and transcriptomic data^16^.

### Modality-specific, complementary subspaces

Each modality will likely have a modality-specific subspace containing information unique to that modality and unpredictable by the other. Although this property confounds applications requiring a shared subspace, it enables other applications because the fusion of two modalities should increase the overall information content, and therefore predictive power, of a profiling dataset.

Data modality fusion and integration techniques are an active area of research in machine learning ^4^ and could potentially yield a superior representation of samples for many different biological profiling tasks on datasets where multiple profiling modalities are available. For example, predicting assay activity might be more successful using information about the impact of that compound on cells’ mRNA levels and morphology, rather than either data source alone ^1^. Likewise, predicting the function of a gene based on similarities to other genes’ profiles might be more successful using both data types.

### Application 1: Cross-modality predictions

As a baseline for finding the correspondence between gene expression and morphology and predicting one from the other, we modeled the relationship using a regression model in which the mRNA level of each landmark gene in the gene expression profile can be estimated as a function of all the morphological features in the Cell Painting profile, *y_l_* = *f*(*X_cp_*) + *e_l_*; in which *y_l_* is a *P*-dimensional (*P*, 1) vector of expression levels for the landmark gene *l* across all the *P* perturbations in a dataset and is the *P*×*F*-dimensional (*P, F*) whole morphological data matrix representing all *F* morphological changes/features across all the *P* perturbations. We use Lasso as a baseline linear model and multilayer perceptron (MLP) as a baseline nonlinear model for the regression problem.

Some datasets showed excellent accuracy in predicting some mRNA levels from morphology data (and vice versa), with MLP yielding superior results to Lasso (Figure 2a and b). Machine learning methods that can improve upon these benchmarks would be very useful to the biomedical community. Two of the datasets (LUAD and LINCS) have a markedly higher performance than the other two (TAORF and CDRP-bio), which suggests a likely poorer data quality or poorer alignment of the modalities in the latter two. Given LUAD and LINCS are both using A549 cells, it is also possible that the transcription-morphology link is cell-line-dependent, and that it is stronger in A549 for some reason; however, it seems even more likely to us that the differences in performance relate to differences in technical quality of the data. Likewise, further preprocessing and denoising techniques such as batch effect corrections to improve alignment are another target for future machine learning research. In addition to alignment across modalities, alignment across different datasets is also necessary to translate the prediction model across different datasets. Application of a model trained on each of the highest performing datasets and tested on the other one (LUAD and LINCS) indicates poor translatability of the models across datasets (Extended Data Fig. 1). Improving model generalizability across datasets requires methods specifically designed to correct for technical variations and batch effects in the bulk level information of the data types presented in this resource.

**Figure 2.**
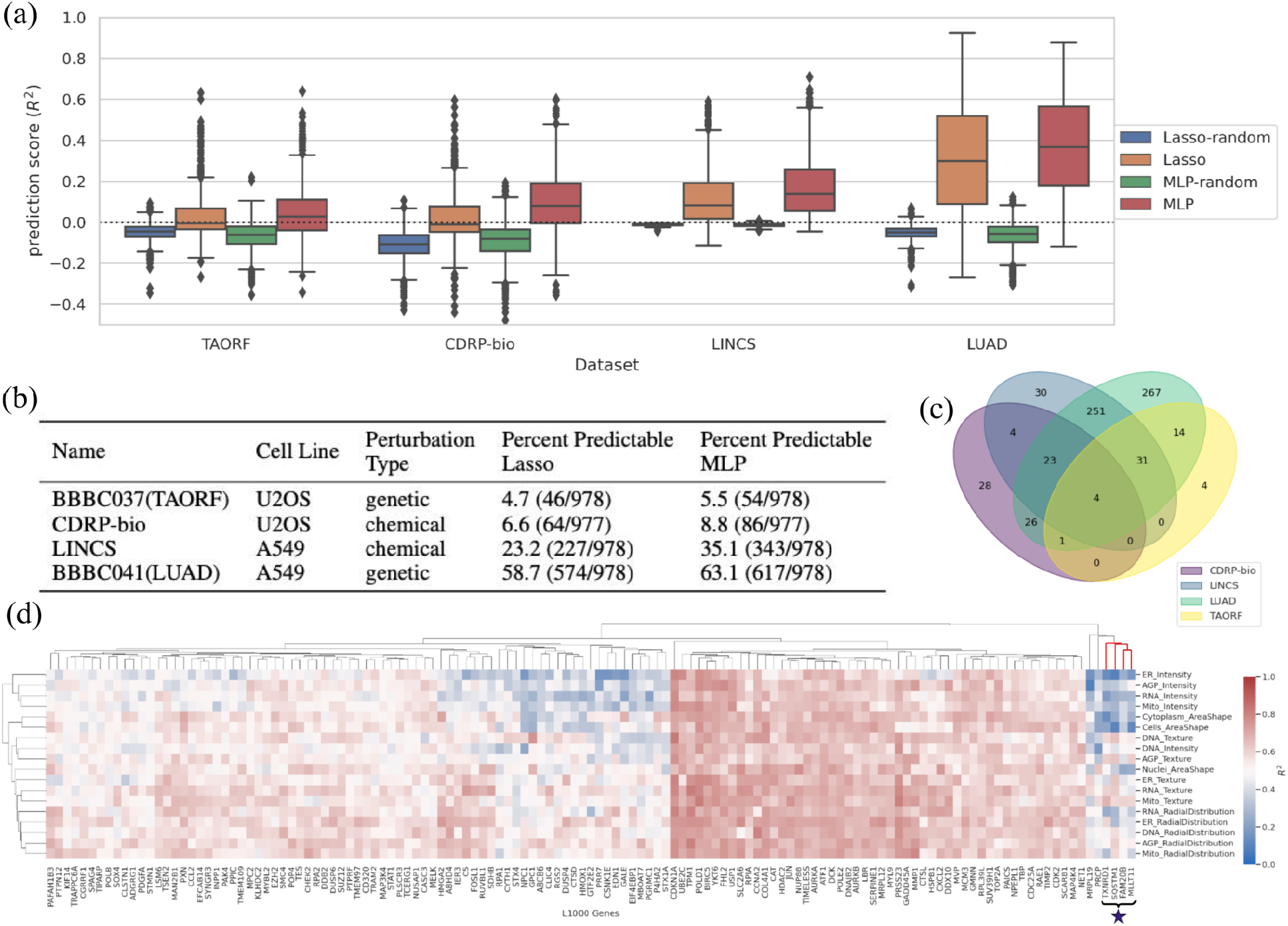
An application using the shared subspace: cross-modality predictions from CP to GE. (a) Distribution of *R*^2^ prediction scores for all landmark genes for each Lasso and MLP model, grouped for each dataset. Many genes are well-predicted, especially using MLP. The random shuffle distributions – where the outputs are shuffled in each iteration– serve as negative controls. Negative *R*^2^ values indicate that the prediction is worse than simply computing the mean of the output, and therefore all *R*^2^ <0 can be considered equally bad (the model does not generalize at all). The y-axis is trimmed at −0.5 for clarity. Distributions are presented as boxplots, with center line being median, box limits being upper and lower quartiles and whiskers being 1.5× interquartile range; n=978 landmark genes (977 for CDRP-bio dataset). (b) The proportion of genes that are well-predicted (*R*^2^ > (*t*_99*th*_ + 0. 2); see Online Methods) are reported as Percent Predictable for each dataset. (c) The overlap of genes predictable by the MLP model (*R*^2^ > (*t*_99*th*_ + 0.2)) are shown across the four datasets; 59 are well-predicted in at least three of the all four datasets. (d) Example of interpretable maps showing the connection between the expression of each landmark gene and the activation of each category of morphological features in the LUAD dataset using the MLP model: each point on the heatmap shows the predictive power of a group of morphological features (on the y-axis) for predicting expression level of a landmark gene (on the x-axis). “Predictive power” here means the *R*^2^ scores generated by limiting the prediction to all the features in the y axis group. The cluster marked with a star is discussed in the main text and explored in Extended Data 4. The heatmap is limited to 131 genes with *R*^2^ > 0.6 scores according to any of the morphological groups (on the y-axis). The complete version is provided in the GitHub repository as an xlsx file (https://github.com/carpenterlab/2021_Haghighi_submitted/blob/main/results/SingleGenePred_cpCategoryMap/cat_scores_maps.xlsx) that can be loaded into Morpheus ^21^ or Python for further exploration.

The shared information in the two modalities can be used in other ways. We can identify the overlap in landmark genes that are highly predictable according to one or more datasets (Figure 2c); 59 landmark genes are well-predicted in at least three of the four datasets. For the LUAD dataset (which has the highest cross-modal predictability) we identified the gene families for highly predictable genes (Extended Data Table 2). Over-Representation analysis of LUAD’s highly predictable gene set (relative to the L1000 background gene set) reveals that many over-represented categories relate to components stained in the Cell Painting assay, such as DNA and actin (Extended Data Fig. 3).

Finally, we examined prediction scores for each category of image-based feature in the experiment, to aid in understanding which features underlie prediction of which genes’ mRNA levels. To do this, we first sorted Cell Painting features into four categories (*intensity, texture, radial distribution, and shape*) and the five fluorescence channels (*DNA, RNA, ER, AGP, Mito*), then calculated and displayed feature-group-specific prediction scores as a hierarchically-clustered heatmap of median (over k-folds) prediction scores (Figure 2d). In this view, genes with strong red columns are predicted using many of the morphological categories of features, indicating that the genes are associated with widespread morphological changes; several of these are cell cycle-related, which is known to impact morphology dramatically. Others are more selective, such as the cluster of genes including TXNRD1, SQSTM1, FAM20B, and MLLT11, whose mRNA levels are strongly predictable by mitochondrial texture features (marked with a star, Figure 2d). Several of these genes have functional annotations relating to mitochondria, and cells that are predicted to have (and actually do have) high levels of these four genes’ mRNA all are associated with visible changes in the staining for mitochondria (Extended Data Fig. 4).

To more generally inspect if the GE-CP relationships observed in Figure 2d are consistent with the known biological functions of the L1000 landmark genes, we performed a Gene Ontology (GO) terms search analysis (see Online Methods). We wondered whether landmark genes that are highly predictable by morphological features in each specific Cell Painting channel are more likely to have Gene Ontology (GO) annotations related to that channel compared to the rest of CP channels; this was generally not the case (Extended Data Table 5; see Online Methods); this is consistent with most predictable genes showing signal across all categories of features rather than being strongly channel-specific (Figure 2d). We also wondered whether landmark genes that are more predictable than other genes are more likely to have functions associated with the particular stains in the Cell Painting assay. Indeed, among the set of 59 highly predictable genes (in at least three of the four datasets) we saw an increased chance of annotations relating to the cellular components and organelles stained in the assay (Extended Data Table 6). That said, many highly predictable genes were associated with no such terms, indicating that the assay probes biological impact beyond the particular labeled components, or that the genes have unannotated functions.

Prediction can be run in the other direction as well, i.e. each morphological feature can also be estimated using the 978 landmark genes as *y_f_* = *f*(*X_ge_*) + *e_l_*; in which *y_f_* is a *P*-dimensional (*P*, 1) vector of measurements for feature *f* across all the *P* perturbations in a dataset and *X_ge_* is the whole gene expression data matrix (*P, L*) representing all *L* landmark genes measurements across all the *P* perturbations. We find a large portion of morphological features to be highly predictable especially for the LUAD and LINCS datasets (Figure 3a). Grouping highly predictable morphological features according to all the datasets reveals that they fall mainly in the radial distribution and texture features categories across all the channels (Figure 3b). We also provide a Jupyter notebook for exploring the list of top connections between any gene or morphological feature of interest (https://github.com/carpenterlab/2021_Haghighi_submitted/blob/main/3-exploreTheLink.ipynb). Users can input an L1000 landmark gene and get the list of top morphological features involved in the prediction of the input feature along with their importance score. Likewise, one can query a morphological feature to find the landmark genes whose mRNA levels are predictive. For example, the morphological feature “Cells_Texture_InfoMeas1_RNA_3_0” relies on the levels of many genes in its prediction, including several known to be involved in mRNA processing (Figure 3c).

**Figure 3.**
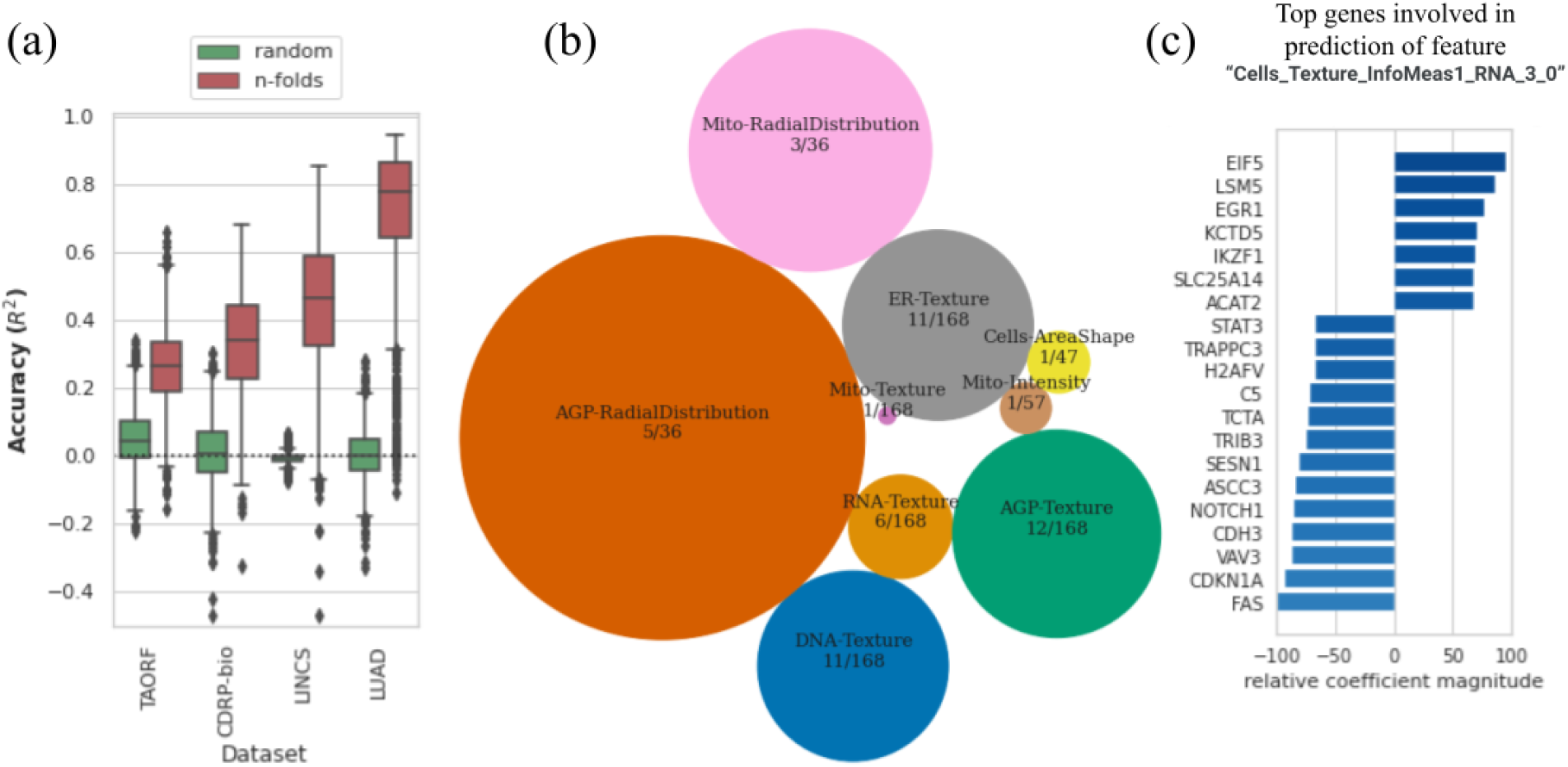
Cross-modality predictions from GE to CP. (a) Distribution of *R*^2^ prediction scores for all morphological features using the MLP model (orange) along with the random shuffle – where the output is shuffled in each iteration – serve as negative controls (green), for each dataset.The y-axis is trimmed at −0.5 for clarity. Distributions are presented as boxplots, with center line being median, box limits being upper and lower quartiles and whiskers being 1.5× interquartile range; the number of points or number of CP features varies among datasets; n=1569 (TAORF), n=1570 (CDRP-bio), n=1670 (LINCS), n=1569 (LUAD). (b) Categories of features with the highest percentage of predictable CP features using GE profiles (median *R*^2^ score across all datasets is more than 0.6). The sizes of circles are proportional to the percentage of highly-predictable features in each category. The number of features in each category over the total number of morphological features in that category are also shown for each circle. (c) Example output of exploratory scripts available to researchers to see what are the most relevant genes to a given morphological feature of interest (and vice versa). The x-axis (relative coefficient magnitude) indicates the relative importance of each feature as the percentage of the strongest feature component (here it translates to the most important landmark gene) involved in the prediction of the morphological feature under exploration. The absolute value and sign of this metric corresponds to the level of importance and direction of the linear relationship respectively. A description of each morphological feature extracted by CellProfiler software is available at: https://github.com/carpenterlab/2016_bray_natprot/wiki/What-do-Cell-Painting-features-mean%3F

### Application 2: Integrating gene expression and morphology

Discerning how a compound works is a major bottleneck in drug discovery. The task is called mechanism of action (MoA) determination, and the goal is to determine the mechanism by which the drug impacts the biological system. Existing methods are often resource- and time-intensive, with a low success rate. As a result, few strategies have been tested systematically across a diverse set of drugs; most inherently only work on a subset of drug or target types, such that multiple methods are usually pursued simultaneously to generate a hypothesis for further testing ^22^. One promising method to predict mechanisms of action is to collect a profile from cells and attempt to match it to a library of profiles gathered from other chemical perturbations: a match, or close similarity, can be helpful if the compound the query matches is already well-known. Likewise, a match to a genetic perturbation means that the gene, or another gene in the same pathway, is a possible target of the query compound ^23^.

Several studies have reported success predicting the mechanism of action of compounds using gene expression or cell morphology data individually ^24–27^ but none of these integrated the two data types to test for improved predictive ability in a supervised or unsupervised setting. We therefore provide here the first benchmark for this, using the two chemical perturbation datasets in our set, CDRP-bio and LINCS. The discovery that many genes could not be well-predicted based on morphology (and vice versa) in Application 1 lends some support for the idea that the two modalities might carry complementary information.

In the unsupervised setting, we test how the compounds cluster together by their MoA class, in feature spaces of each modality alone, and in the integrated space of both modalities, using several state of the art modality integration methods^28^. Clustering of perturbations using each CP and GE modality alone shows CP outperforms GE in this MoA ground-truth retrieval task in both compound datasets. We observe that although most of the integration methods increase cluster retrieval performance in the integrated space compared to the GE space, only Regularized generalized canonical correlation analysis (RGCCA)^29^ improves the performance over the CP space alone (Figure 4a).

**Figure 4.**
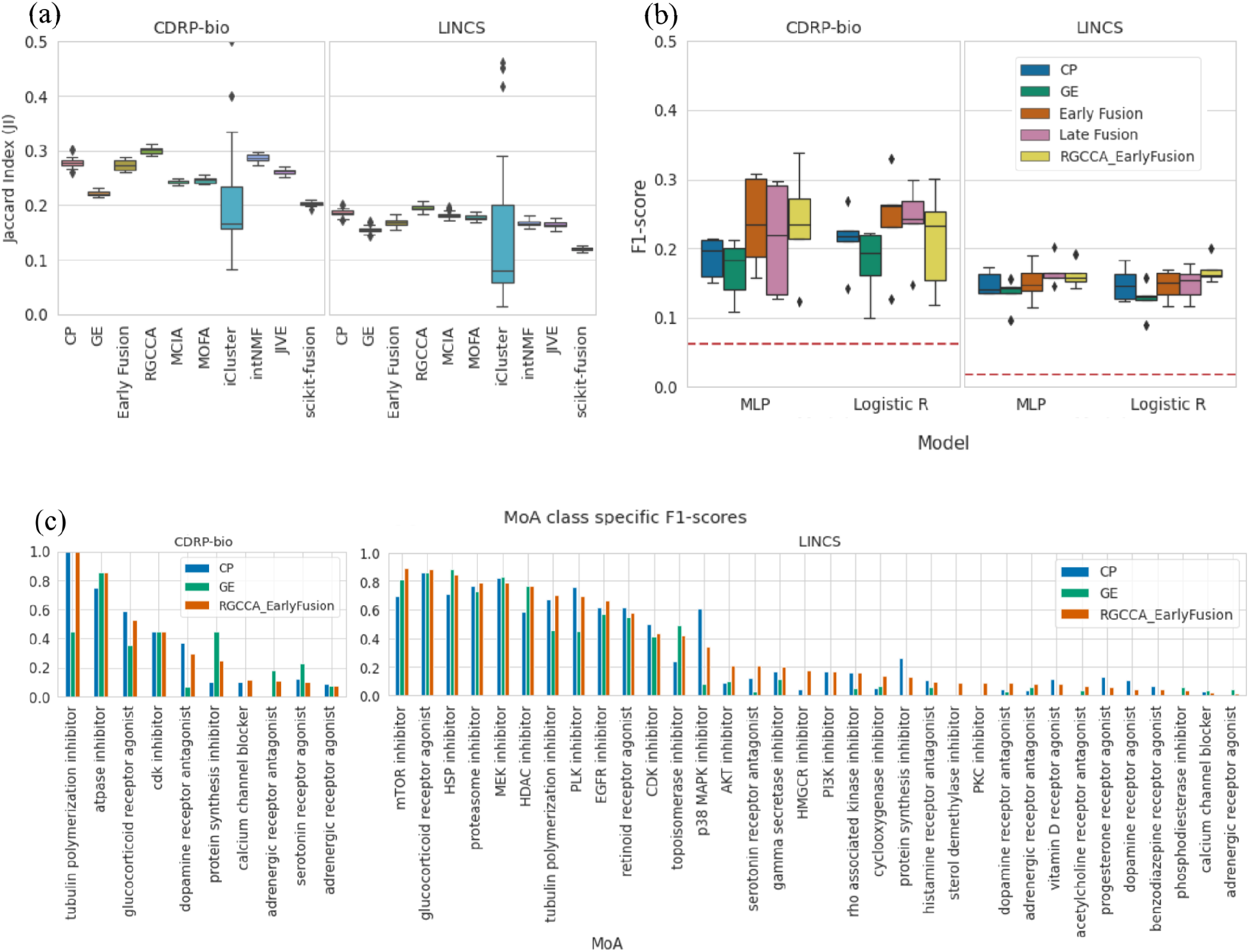
Utilizing Complementary Information: Data Integration for MoA cluster retrieval and class prediction in compound datasets. An application using the complementary subspaces: integrating multimodal data for mechanism of action (MoA) unsupervised clustering retrieval (a) and supervised prediction (b): (a) Benchmarking of data integration methods on the task of clustering compounds by their MoA categories. Distribution of the Jaccard Indices (one per MOA class; higher is better) computed between the clusters identified by the different integration methods^28^ and the ground-truth MoA clusters. Regularized Generalized Canonical Correlation Analysis (RGCCA) improves MoA retrieval for both CDRP-bio and LINCS datasets. Distributions are presented as boxplots, with the center line being median, box limits being upper and lower quartiles and whiskers being 1.5× interquartile range; n=16 (CDRP-bio), n=57 (LINCS) (b) MoA classification of the two compound datasets (CDRP-bio and LINCS) using gene expression, morphology and their integration to predict the mechanism of action of compounds. Classification performance (weighted F1-score) for the multilayer perceptron (MLP) and Logistic Regression classifiers using each data modality alone, the two early and late fusion strategies explained in the main text, and the early fusion of modalities after application of RGCCA on the feature space of both modalities. Chance-level predictions for each dataset are shown as a horizontal red line on each dataset plot. Distributions are presented as boxplots, with the center line being median, box limits being upper and lower quartiles and whiskers being 1.5× interquartile range; n=k=5. (c) Class-specific F1-scores are shown based on the MLP model for 16 MoA categories of CDRP-bio (left, where the 4 out of 16 MoA categories that resulted in zero F1-scores after fusion are excluded) and for LINCS (right, where the 23 out of 57 MoA categories that resulted in zero F1-scores after fusion are excluded).

In the supervised setting, using logistic regression and multilayer perceptron (MLP) classifiers as the baseline models, we applied each for predicting MoA labels using each modality of data independently, using a standard k-fold (k=5) cross-validation on a filtered subset of compounds. CP profiles resulted in higher MoA prediction performance compared to GE profiles for each of the datasets (Figure 4b). Next, we performed the MoA prediction task on two integrated spaces: 1) trivial representation-level concatenation of profiles from the two modalities (shown as early fusion) and 2) representation-level concatenation of profiles from the two modalities in the RGCCA space (shown as RGCCA_EarlyFusion). We also performed decision-level integration of modalities for the MoA prediction task (shown as late fusion), i.e. class probabilities output by classifiers trained on each modality separately are averaged prior to the final MoA prediction.

All three integration strategies gave relatively comparable performance in predicting MoA across the two datasets and two model types, with small average improvements upon the performance of the better-performing modality (Figure 4b), highlighting the need for developing data fusion methods that better leverage the complementarity of the modalities.

Exploring MoA-class-specific F1-scores for the integrated modalities reveals high variation in class-specific prediction results (Figure 4c). As already seen more generally, the integration of modalities does not always increase the performance of the MoA prediction task over the higher performing modality alone for individual MoA categories.

## Discussion and Limitations

We provide the research community a collection of multi-modal profiling datasets with gene expression and morphology readouts, representing two cell types and two perturbation types (genetic and chemical). We define useful biological applications for this data in two categories: those using the shared information and those using modality-specific, complementary information. We provide the data, code, metrics, and benchmark results for one application in each category.

The results demonstrate that gene expression and morphology profiles contain useful overlapping and distinct information about cell state. We were pleased to find that many mRNAs are predictable by cell morphology and vice versa, under the conditions of these high-throughput assays. Similarly, we found that morphology captures information beyond that seen in an mRNA profile; that is, the two modalities contain unique information and we identified which compounds’ mechanisms are better captured by each. Although some scientists speculated that it is impossible for cells to show a morphological change without mRNA profiles changing, whether as a cause or consequence, we find this is not the case. We made a number of observations of new biology, such as which genes’ mRNA levels are predicted by which particular morphological features (and vice versa). Finally, we discovered that the Cell Painting assay information can predict the mRNA levels of genes not clearly linked to the stains in the assay, pointing to its ability to capture broad biological impact.

The results also demonstrate that these applications are challenging enough to provide room for improvement. For example, the variation in the performance for prediction tasks across different datasets shows the necessity of machine learning techniques to further filter and preprocess the profiles (e.g. to correct batch effects, including those resulting from the position of wells on a plate^30^) to improve performance. Such techniques might also sufficiently align the four datasets with each other, to explore generalized, dataset-independent models. Nevertheless, we note that we do not expect anywhere close to 100% accuracy for either application. For prediction across the two modalities, we do not expect the modalities to be completely overlapping in their shared information. Furthermore, we note that ground truth in this prediction task is defined only by the available experimental gene expression and cell morphology data, which is subject to technical variation and error and therefore is not absolute truth. In the case of MoA prediction, the application is “notoriously challenging” and low percentage success rates are expected for any single assay; most commonly several strategies are used to determine the mechanism of action ^31^. In addition, the ground truth is based on imperfect human knowledge.

There are multiple additional limitations for the presented datasets, aside from their data quality as already noted. The number of gene perturbations captured in these datasets number a few hundred whereas there are roughly 21,000 genes in the genome and numerous variations within each, which could be overexpressed or knocked down. Likewise, a few thousand compounds are tested here but pharmaceutical companies often have collections of compounds numbering in the millions. The only limitation for expanding these datasets are the financial resources to carry out the experiments. In terms of limitations of the assays themselves, the gene expression profiles are captured by the L1000 assay, which is thought to capture 82% of full transcriptome variation ^7^, and the Cell Painting assay includes only six stains, which is insufficient to capture the localization and morphological variation of all cellular components. Finally, the cell types are commonly used historical lines derived from two white patients, one male (A549) and one female (U2OS). Therefore, conclusions from this data may only hold true for the demographics or genomics of those persons and not broader groups. They were chosen because the lines are both well-suited for microscopy and they offer the advantage of connecting to extensive prior studies and datasets using them.

Despite these limitations, these datasets may be used to pursue many other applications of profiling in biology, as well as methods development. The complementary information used here for MoA prediction can be used for any profiling application; there are more than a dozen that can impact basic biology discovery and the development of novel therapeutics^32^. Each application can also be validated in different ways. For example, the prediction task might be extended to more complex systems, such as human tissue samples where appropriate stains have been used, although such samples are more difficult to procure, and assessing adjacent tissue slices may introduce variation not present in the cultured cell lines used in this study. In the future, multimodal profiles at the single-cell level may become widely available. In the presented datasets, single-cell information exists in one modality (images) but not in the other modality (mRNA). Therefore, the variations in one cannot be explained by the other, as we have a distribution in one space (images) and point estimates in the other space (mRNA). Although still very rare, small, and labor-intensive to create, data sets with both gene expression and morphology at single-cell resolution are beginning to become available via in situ RNA-seq methods and could accelerate the field of multi-modal biological data analysis.

## Data Availability

Preprocessed profiles that are augmented with gene and compound annotation are available through the *Registry of Open Data on AWS* on a public S3 bucket at no cost and no need for registration. Documentation on the folder structure, dataset details and instructions for accessing the data are available at http://broad.io/rosetta. Source datasets are described and referenced in Supplementary 1.

## Code Availability

Source code to reproduce and build upon the presented results is available at http://broad.io/rosetta. We license the source code as BSD 3-Clause, and license the data, results, and figures as CC0 1.0.

## Acknowledgements

We thank all the researchers who created and shared the data, who are mentioned in their respective publications cited in the paper.

Funding was provided by grants 2018-183451 and 2020-225720 to AEC from the Chan Zuckerberg Initiative DAF, an advised fund of the Silicon Valley Community Foundation and the National Institutes of Health NIGMS (R35 GM122547 to AEC).

## Author information

M.H., S.S., B.A.C, and A.E.C. all contributed to drafting the manuscript and designing the research. J.C. initiated the project and performed early explorations of the LUAD dataset. M.H. analyzed and explored the data with inputs from the other coauthors.

## Online Methods

### Dataset preprocessing

We gathered four available datasets that had both Cell Painting morphological (CP) and L1000 gene expression (GE) profiles, preprocessed the data from different sources and in different formats in a unified .csv format, and made the data publicly available at amazon s3 bucket:

~~~
s3://cellpainting-gallery/cpg0003-rosetta/broad/workspace/preprocessed_data/
~~~

The rows of each csv file are replicate-level (i..e well-level) profiles, augmented with metadata available for that well.

#### Cell Painting and L1000 Profiles

Single-cell morphological (CP) profiles were created using CellProfiler software and processed to form aggregated replicate profiles using the R [cytominer](https://github.com/cytomining/cytominer/blob/master/vignettes/cytominer-pipeline.Rmd) package (https://github.com/cytomining/cytominer).

We made the following three types of profiles available:

- Aggregated profiles, which are the average of single-cell profiles in each replicate well. (*replicate_level_cp_augmented.csv.gz*)
- Normalized profiles, which are the z-scored aggregated profiles, where the scores are calculated using the distribution of negative controls as the reference. (*replicate_level_cp_normalized.csv.gz*)
- Normalized variable-selected, which are normalized profiles with features selection applied. (*replicate_level_cp_normalized_variable_selected.csv.gz*)

For L1000, we use the previously processed 978 “landmark” genes as our input features The complete processing details are provided in ^7^. The L1000 landmark genes in the CDRP dataset are different from the landmark genes in the other datasets, with an overlap of n=785 (80%). The CDRP dataset was acquired using the so-called *“delta prime”* probe set (n=977). Subsequent datasets (LUAD, TAORF, LINCS) were acquired using the so-called “*epsilon*” probe set (n=978)^33^.

The 20% of the delta prime landmark genes that are absent in epsilon can be inferred using the epsilon landmark genes ^7^.

To simplify our analysis, we did not perform this inference, and instead only used the landmark genes available for each dataset. When combining CDRP with other datasets, we used the intersection of the two probe sets.

#### Data processing for Analysis

We have used treatment-level profiles for both the gene expression (GE, using L1000) and morphology (CP, using Cell Painting) modalities for the analysis presented, although replicate-level profiles are provided and could be used instead in other formulations of the problem to create more advanced models.

Treatment-level profiles are the average of replicate-level profiles. For CP, replicate-level profiles are the average of single-cell measurements of cells from that replicate well. For GE, replicate-level profiles are simply the bulk gene expression profile for that replicate well.

We standardized replicate-level profiles per plate to have zero mean and unit variance before averaging them to form treatment-level profiles.

Note that, aside from some image segmentation parameters in the CellProfiler pipeline which are adjusted for each cell type based on its baseline morphology, the computational pipeline for data processing and analysis were identical regardless of the cell type in the experiment.

#### Measuring quality of data points for subsequent analysis

We use treatment-level profiles for all the analysis that follows. In the rest of the text, unless indicated otherwise, “data points” refer to treatment-level profiles. The specific transformation of the treatment-level profiles (such as *normalized_variable_selected*) is clarified when necessary.

There are inherent differences in the biological design (type of perturbation, cell line used, and time point of exposure to perturbation) and experimental parameters (different instrumentation, reagent batches, and personnel running the experiments creating distinct technical artifacts such as batch effects) differences in the datasets. Consistency of profiles of a single treatment across different batches of experiment is considered a measure of data quality. We check this consistency as follows. After standardization of the replicate-level profiles per plate, we calculate the Pearson correlation coefficient between each pair of replicate-level profiles for the same perturbation. The distribution of these coefficients for each dataset and modality are illustrated in Supplementary Figure 1 shown as red curves. The corresponding blue curve to each red curve is the null distribution showing the correlation coefficient between pairs of profiles that belong to different perturbations. The non-zero dotted vertical line to the right shows the 90th percentile of the null distribution. We consider the perturbations that have an average replicate correlation more than the 90th percentile of the null distribution as high quality data points for subsequent analysis.

We note a source of systematic error present in all datasets that may affect replicability metrics: for nearly every treatment, all its replicates occur at the same well position on the plate (because replicates in such high-throughput experiments are created by replicating the entire, and exact same, plate layout, for logistical reasons). The location of the well on the plate can impact the cells in the well. For example, wells on the edge are more likely to dry slightly, impacting cell morphology. This effect – the impact of an experiment covariate on the readout of the assay – can inflate replicability quality metrics. In our experience, well-position effects tend to be more pronounced in Cell Painting than L1000, and therefore the observed differences in data quality – as reported in Supplementary 2 – can be a function of this batch effect. As we note in the discussion, correcting for batch effects could improve the prediction tasks discussed in this paper, and also make such comparisons of data quality more reliable.

#### Filtering data points

To remove noisy data points from the analysis, we used two filtering strategies for each shared subspace and data integration analysis. For cross-modality prediction experiments, we used the intersection of higher quality data points according to both modalities. For the analysis for data integration, we used data points that are higher quality (i.e. > 90th percentile of the null distribution, as defined above) in at least one of the modalities. Definition of higher quality data points is given in the previous section. A comprehensive description of the data sizes in each modality, number of overlapping perturbations across both modalities, size of intersection and union sets of higher quality data points across both modalities are given in Supplementary 1 and highlights are summarized in Supplementary Table 1.

One of the chemical datasets (CDRP-BBBC047-Bray) has a subset of compounds that are known to be bioactive. We refer to this subset as CDRP-bio-BBBC036-Bray and report the details independently for this dataset in Supplementary 1 and 2. We only use CDRP-bio and not the full CDRP set for the analysis in this paper. We did so because we believe that the quality of CDRP is insufficient for either of these analyses presented given that very few data points remain after filtering for replicate reproducibility across both modalities (see Supplementary Figure 1).

### Cross modality Predictions

For prediction of each single landmark gene using CP profiles or each single morphological feature using GE profiles, we used two regression models of:

#### CP to GE

*y_l_* = *f*(*X_cp_*) + *e_l_* in which *y_l_* is a vector of expression levels for the landmark gene *l* across all the perturbations in a dataset and *X_cp_* is the whole morphological data matrix where each row is a treatment-level Cell Painting profile. For this prediction direction, we have used the so-called *“normalized variable-selected”* treatment-level Cell Painting profiles, which results in 601 features for CDRP-bio dataset, 291 features for LUAD dataset, 63 features for TA-ORF and 119 features for the LINCS dataset. The variable selection step removes features with near zero variance as well as reduces redundancy in the feature set (ensuring that no pair of features have a Pearson correlation coefficient > 0.9).

#### GE to CP

*y* = *f*(*X_ge_*) + *e_l_*; in which *y_f_* is a vector of morphological feature *l* across all the perturbations in a dataset and *X_ge_* is the whole gene expression data matrix where each row is a treatment-level L1000 profile. For this prediction direction, we have not performed any dimensionality reduction on the GE data.

For each prediction direction (CP to GE, GE to CP) and each baseline linear (Lasso) and nonlinear (MLP) model for this regression problem, we use the coefficient of determination (*R*^2^) and nested *k*-fold cross-validation over the data points for evaluating the prediction model performance. Therefore, for each landmark gene (for CP to GE) or each morphological feature (for GE to CP), we can form a distribution of *k*, *R*^2^ values. We also shuffle the vector *y_l_* for each gene *l* across all the data points and apply the same cross-validation procedure to form a null distribution for each gene. The same procedure on *y_f_* will result in the null distribution for each morphological feature. Model parameters are selected using grid-search and cross-validation on each training set for each of the *k* test folds.

In the Supplementary 4, the median prediction scores of each model for each landmark gene for each dataset and according to each model is presented. Distribution of MLP model prediction scores for the 50 landmark genes with the highest median scores in each dataset is shown at Supplementary 3.

##### Percent Predictable

Percent predictable is defined as the percentage of landmark genes which have a median of *R*^2^ predictability score more than a defined threshold. The threshold is based on the null distribution of predictability scores for each dataset. The dataset-specific null is formed using medians of single gene null distributions. We take the 99th percentile of this null distribution plus a 0.2 margin (*t*_99*th*_ + 0.2) as the threshold for calling a gene “predictable”. We reported the *Percent Predictable* values for each dataset in the table in Figure 2b.

### Modality integration

For the analysis for MoA prediction, we used the data points that had high quality according to their replicate-level profiles (i.e. > 90th percentile of the null distribution; see above) in either modality. Note that in compound datasets, each perturbation is tested at multiple doses and therefore there are multiple data points corresponding to each compound. A data point here is a treatment-level profile corresponding to a dose of a compound.

The LINCS dataset has MoA annotations for 1401 overlapping compounds across two modalities. Every compound is tested at seven different doses, increasing the chances of detecting the expected behavior of the compound at one of them. Each compound can have multiple mechanisms, therefore we have multiple labels for a subset of compounds. The set of labels comprises 478 unique MoAs. There are 568 unique combinations of these labels present in the dataset. We start with the filtered union set, and filter it again to keep MoA classes which have at least 5 data points in their class. Because this process resulted in only one MoA category that was multi-label (i.e. composed of multiple MoAs), we removed this category to simplify the problem as being multi-class but single label i.e. we effectively used only compounds labeled with a single MoA. The filtered set is a set of 1655 data points across 521 compounds in 57 MoA categories.

The CDRP-bio dataset has MoA annotations for 1,327 out of 1,916 overlapping compounds across two modalities. After passing data points from three filters – union higher quality across modalities, available MoA labels, being in an MoA class which have at least five compounds in the set – we get 123 compounds in 16 MoA categories.

#### Unsupervised joint dimensionality reduction of modalities and MoA cluster retrieval

k-means clustering (k= number of MoA classes) was performed on each modality space and integrated spaces using representation-level concatenation of modalities (early fusion) and seven state of the art modality integration methods^28^. Jaccard Index (JI) between k-means clustering results and the MoA annotation labels were used as a measure of ground truth cluster retrieval for this unsupervised clustering task.

#### Supervised MoA Prediction

For the multi-class MoA classification problem, two logistic regression and multilayer perceptron (MLP) classifiers were used as baseline models; we apply each model for predicting MoA labels using each modality of data independently as well as the baselines for integration of the two. We performed stratified nested k-fold cross-validation (k=5) to evaluate the classification performance using the F1-score metric. Note that all doses of a compound should be in the same fold in this data partitioning scheme. Model hyper-parameters were optimized using grid search and cross-validation in each training fold.

Some MoAs have several tens of compounds whereas others have as few as 5, to address this imbalance in the data, for both logistic regression and MLP models, we oversampled data points in each class to match the size of the majority class in the training set. The k-fold cross-validation experiment results in *k* vector of multi-class predictions. We then calculate F1-score for each class independently and average class specific F1-scores within each fold will form *k* F1-scores shown in Figure 4b boxplots.

For representation level integration strategies, we simply concatenate CP and GE profiles in their original spaces (Early Fusion) or projected into RGCCA space (RGCCA_EarlyFusion) to integrate both modalities. On the other hand, Late fusion is at the decision level and averages predicted class probabilities (based on the output of classifiers trained on each modality separately) for making the MoA class decision for test compounds.

### Gene Ontology (GO) terms search analysis

The goal of this analysis was to see if the found CP-GE link is consistent with the known functional characteristic of L1000 landmark genes under study in this work. We used DAVID^34^’s Functional Annotation Tool (2021 Update) to form a table of GO annotation terms for 978 landmark genes in the LUAD, LINCS and TAORF datasets. For 921 DAVID IDs detected, GO categories of *“GOTERM_BP_DIRECT”, “GOTERM_CC_DIRECT”* and *“GOTERM_MF_DIRECT”* were selected and the *Functional Annotations Table* were used as the source file for the following search analysis.

For each landmark gene and each cell painting channel, we searched for all relevant keywords to that channel and formed channel specific annotation columns. On the other hand, mRNA level prediction of landmark genes using specific categories of morphological features were created for LUAD dataset (Figure 2d), we form channel specific gene expression prediction scores by rearrangement of these categories to cell paining channels and taking the maximum prediction score within each channel specific category of features. We then have access to channel specific GO functional annotations and channel specific prediction scores for each of the landmark genes. We discretize prediction scores into three categories of predictability; high(*R*^2^>0.8), medium and low(*R*^2^ <0.1). Using Fisher’s exact test, we calculate Odds Ratio (OR) as a measure of association between a landmark gene being predictable and having channel-specific GO annotations. OR>1 indicates an increased chance of having an organelle annotation (using GO terms) for a highly predictable landmark gene.

## Extended data

### Extended Data 1. Generalizability of the prediction model across datasets

**Extended Data Fig. 1.**
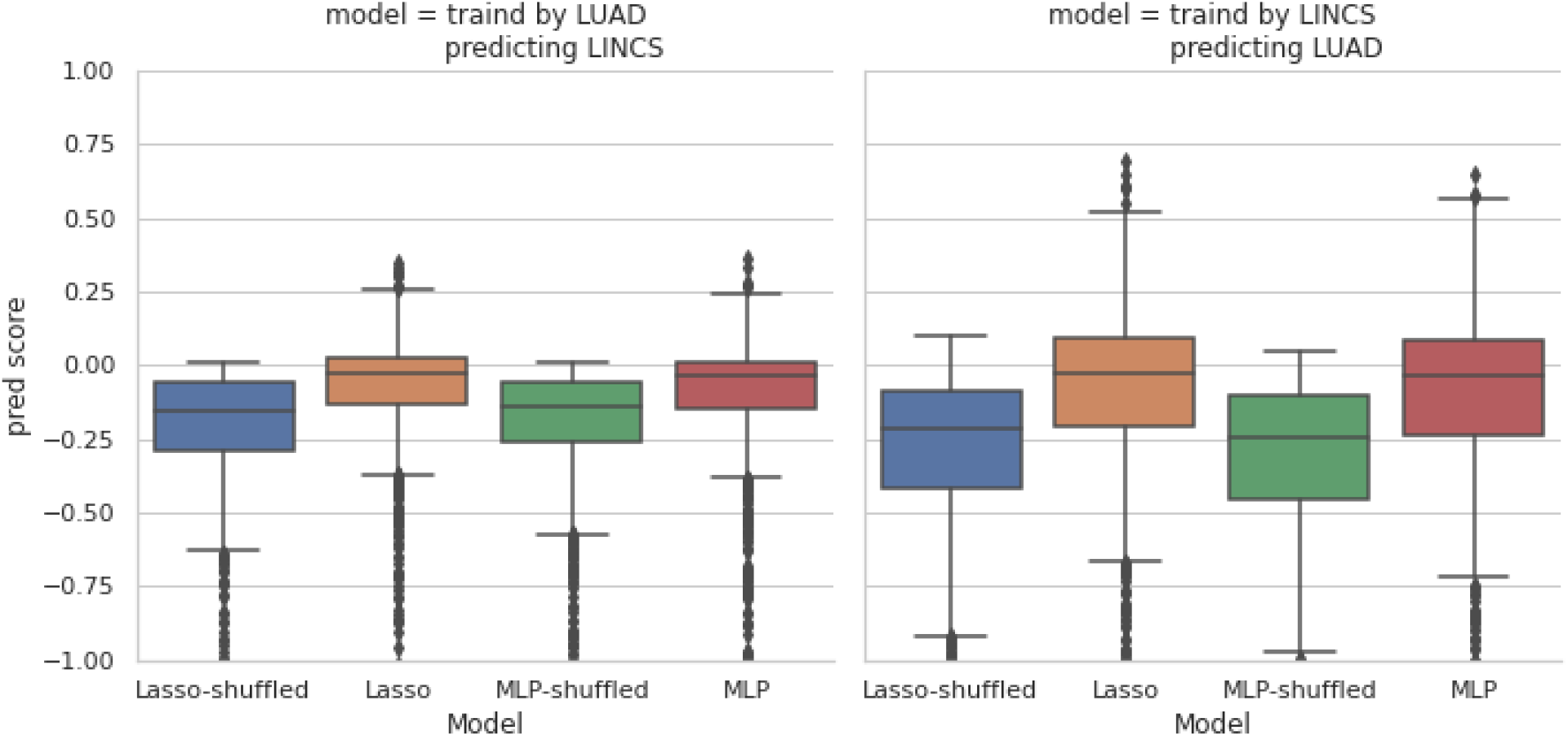
Prediction of each L1000 mRNA level by Cell Painting features in dataset A, using a model trained on dataset B. We have trained Lasso and MLP models on each of LUAD and LINCS datasets and checked the prediction results on the other dataset which was not used in model training. Distribution of *R*^2^ prediction scores for all landmark genes are shown. Comparison of the results here with Figure 2 indicates weakness of the prediction model in generalizability across datasets. This is an indication of dataset-specific technical variations (batch effects) that need exploration of experimental alignment techniques (batch-effect correction), which is an active area of research. We also observe that the model’s prediction power is stronger when the model is trained on the LINCS dataset and tested on the LUAD dataset. This is expected as the LUAD dataset is limited to a narrow set of genes associated with lung adenocarcinoma cancer; however, the LINCS dataset contains a wide variety of compounds with different mechanisms and known phenotypes. The y-axis is trimmed at −1 for clarity. Distributions are presented as boxplots, with center line being median, box limits being upper and lower quartiles and whiskers being 1.5× interquartile range; n=978 landmark genes for each boxplot.

### Extended Data 2. Gene group names for top 100 predictable landmark genes in LUAD dataset

**Extended Data Table. 2.**
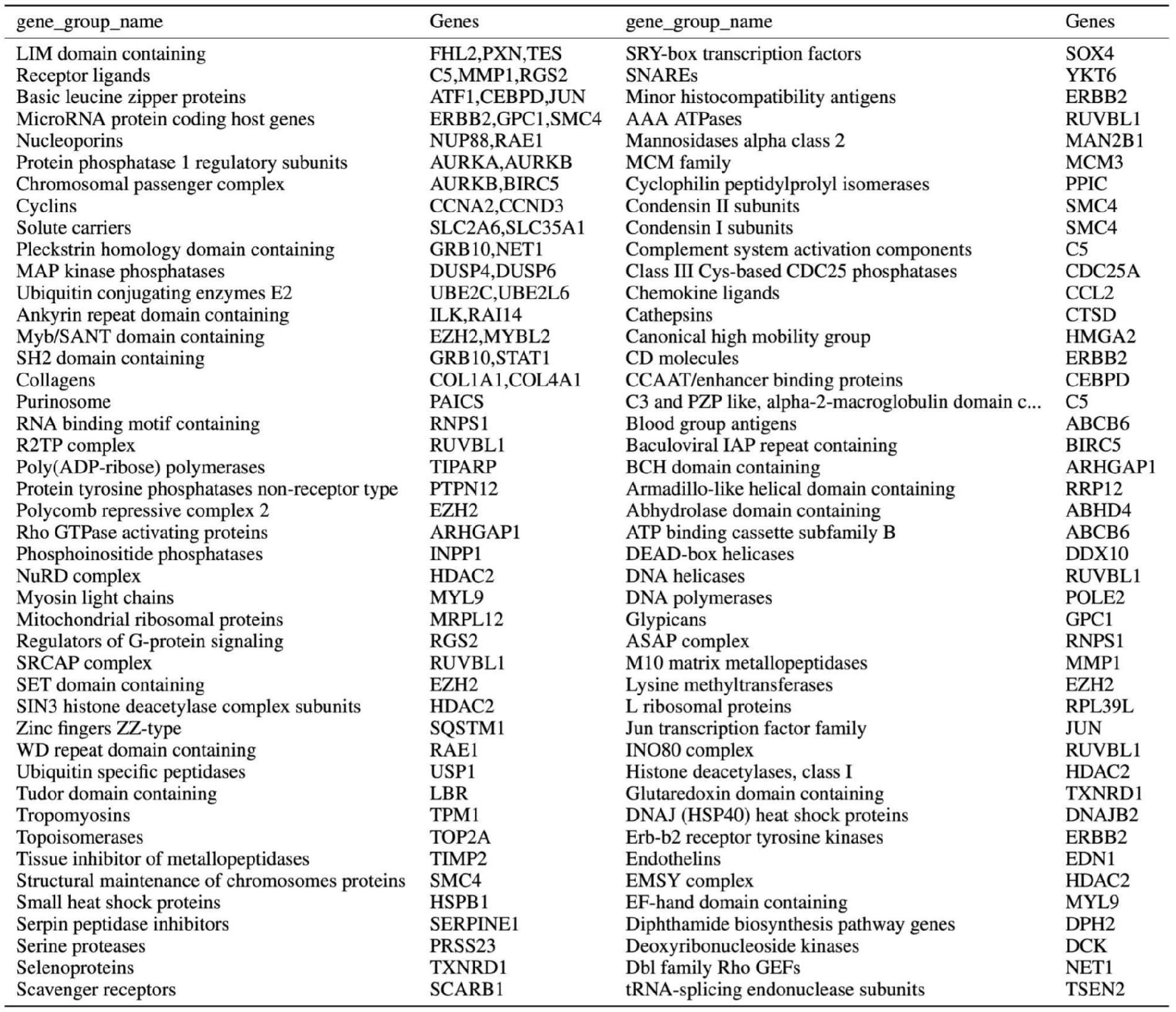
Top 100 predictable landmark genes by MLP model are shown along with their gene group names (based on HGNC Database^41^) for the LUAD dataset, finding a diverse array represented, though we note the perturbations in this experiment included only genes found mutated in lung cancers.

### Extended Data. 3. Over-Representation Analysis (ORA) of highly predictable (top 100) landmark genes in LUAD dataset

**Extended Data Fig. 3.**
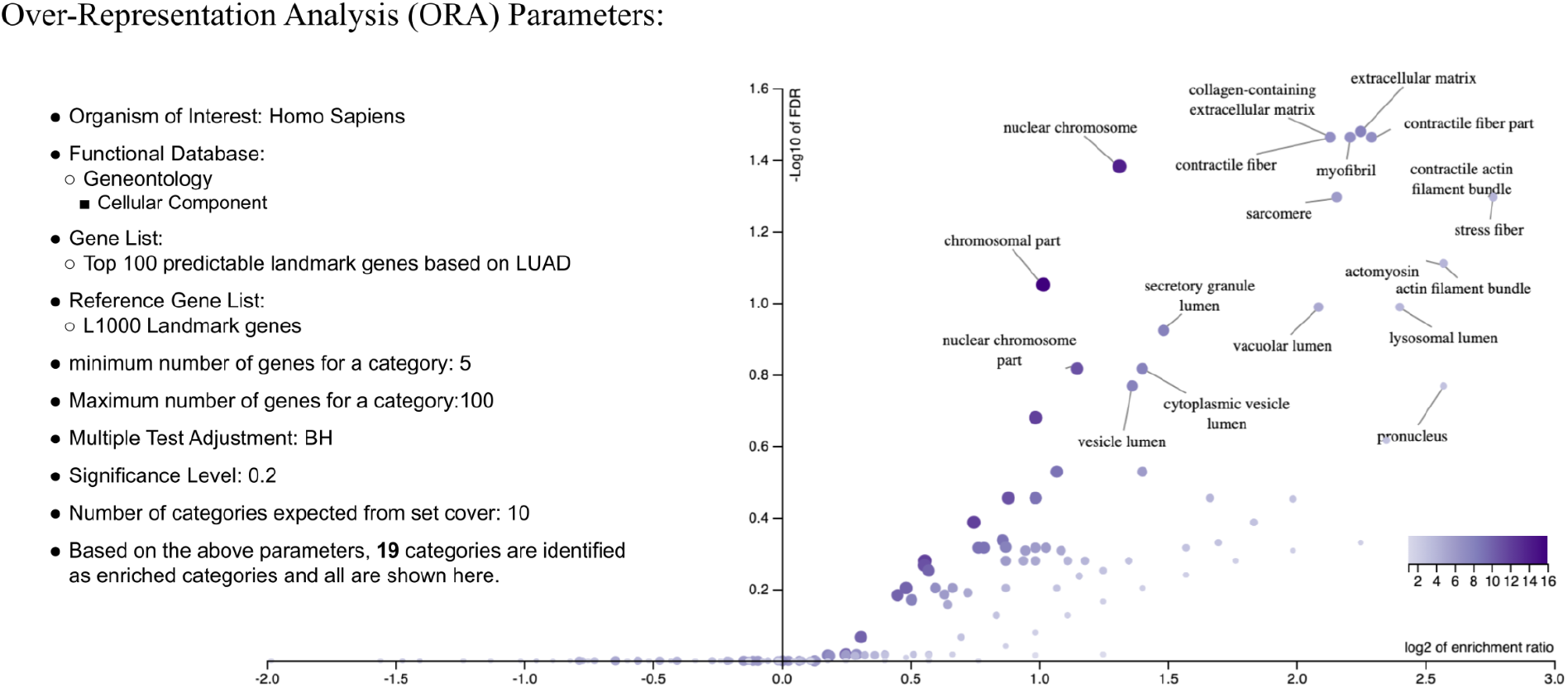
Over-Representation Analysis of top 100 highly predictable landmark genes according to the MLP model applied on the LUAD dataset. ORA analysis was performed by *WebGestalt* analysis toolkit^42^. Nineteen enriched categories (FDR<0.2) are labeled in the volcano plot.

### Extended Data. 4. Visualization of cells in a cluster of landmark genes that are tightly correlated with RNA texture category of morphological features

**Extended Data Fig. 4.**
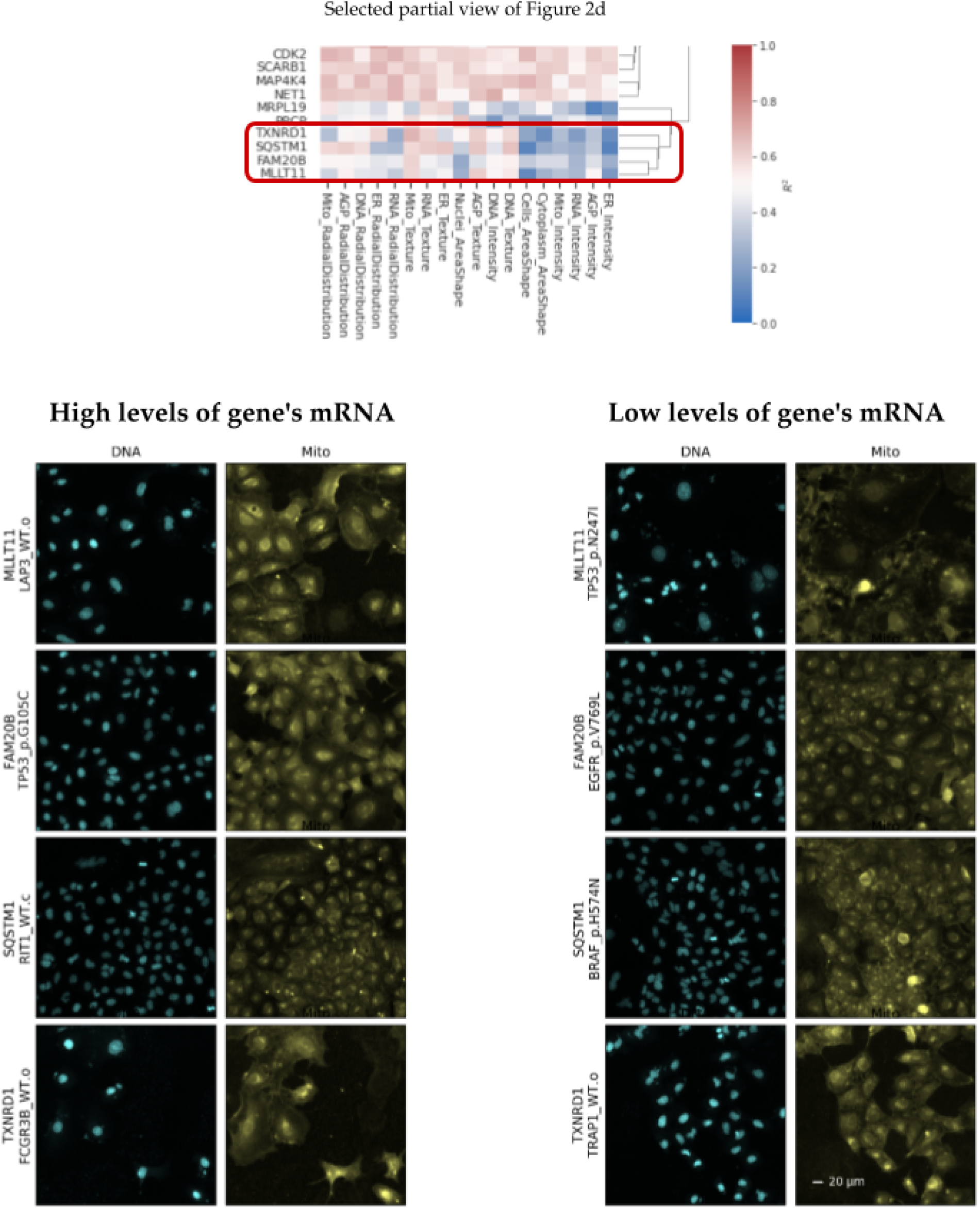
For the cluster of landmark genes shown in the top heatmap, which is a partial snapshot of figure 2d, we have shown example cell images for perturbations that have high and low predicted values for each gene in that cluster. We have filtered perturbations to those that have low prediction errors prior to that selection. We can observe that cells that are predicted to have (and actually do have) high levels of these five genes’ mRNA all are associated with visible changes in the staining for mitochondria, even though only half of these genes already have functional annotations related to the mitochondria.

### Extended Data. 5. Validation of the observed GE-CP relationship by GO-terms search analysis

**Extended Data Table. 5.** Landmark genes highly predictable according to morphological features in each specific Cell Painting channel are more likely to have GO annotation related to that channel compared to the rest of CP channels. For each channel in the rows of the table, the first column shows the Odds Ratio (OR) derived from the Fisher’s exact test for associations between the landmark genes being highly predictable (*R*^2^ >0.6) by CP features in a channel and having GO annotations for that channel. The second column shows the association between the same set of highly predictable genes and having GO annotation for any channel but not the target row channel. Higher values in the first column compared to the second column show that highly predictable genes according to features in a CP channel are more likely to have GO annotations for that channel compared to the rest of the channels. This pattern holds for DNA and ER channels but not for the rest of CP channels. The third and fourth columns show the same associations but for low-predictability genes (*R*^2^<0). Lower values in the third column compared to the fourth column show that non-predictable genes according to features in a CP channel are less likely to have GO annotations for that channel compared to the rest of the channels. This pattern holds for all CP channels except for RNA. The CP channel specific predictability map used for this analysis was derived from the result of the experiment and results presented partially in Figure 2d. As we can observe from the map, usually multiple categories of morphological features contribute to the predictability of a gene, which explains the lack of a simple relationship between a given channel’s predictability and GO term associations presented in this table.

| CP channel | High predictability |  | Low predictability |  |
| --- | --- | --- | --- | --- |
|  | Odds Ratio |  | Odds Ratio |  |
|  | same channel | rest of channels | same channel | rest of channels |
| DNA | 2.188 | 0.643 | 0.720 | 6.953 |
| RNA | 0.952 | 1.004 | 1.375 | 1.165 |
| AGP | 0.851 | 1.042 | 0.741 | 0.974 |
| Mito | 0.578 | 1.381 | 0.896 | 0.946 |
| ER | 1.040 | 0.829 | 0.598 | 1.070 |

### Extended Data. 6. Association between landmark gene predictability and having gene ontology annotations related to Cell-Painting stains

**Extended Data Table. 6.** Landmark genes that are predictable according to at least three of the four datasets (59 genes shown in Figure 2c) are more likely to have GO annotations related to any of the stains in the Cell Painting assay compared to a random subset of landmark genes.

| Stains | Mitochondria | Golgi | Membrane | Cytoskeleton/Actin | Endoplasmic | RNA/Nucleoli | DNA | Any stain | No stain |
| --- | --- | --- | --- | --- | --- | --- | --- | --- | --- |
| Odds Ratio | 1.58 | 0.69 | 1.22 | 1.59 | 1.02 | 0.64 | 1.95 | 2.5 | 0.4 |

