## Supplementary information for "High-Dimensional Gene Expression and Morphology Profiles of Cells across 28,000 Genetic and Chemical Perturbations"

### Supplementary 1. Curated Datasets

**Datasets List**
The details of each dataset, including the type of perturbation, number of perturbations, cell line, and number of replicates, are below. All experiments were performed in multi-well plates (384-well). The plate design varies across the datasets. The design can be visualized for each plate by inspecting any of the CSV files for each dataset (for example, for Cell Painting, *Metadata_Plate* and *Metadata_Well* provide the coordinates, and the rest of the columns with a "*Metadata_*" prefix have details of each perturbation).

**CDRP-BBBC047-Bray-CP [35]- GE [7]:**

The U2OS cell line (ATCC catalog #HTB-96) was used in this chemical perturbation dataset. For CP and GE data, data was collected after 6 and 48 hours of compound treatment, respectively; the plate maps are available at s3://cytodata/datasets/CDRPBIO-BBBC036-Bray/metadata/CDRPBIO-BBBC036-Bray/platemap/. Cell Painting was performed using the Cell Painting v1 protocol [36]. Widefield imaging was performed on an ImageXPress microscope with a 20X objective with 2x2 binning, resulting in 696x520 images. Illumination correction, segmentation, and feature measurement were performed in CellProfiler 2.2.0, per pipelines available at https://github.com/gigascience/paper-bray2017/tree/master/pipelines. L1000 was performed using the protocol described in [37].

There are 30,430 and 21,782 unique compounds for CP and GE datasets, respectively.

For CP dataset, the median number of replicates for each compound in the set is 4 and there are 26,572 replicates of control wells.

For GE dataset, the median number of replicates for each compound in the set is 3 and there are 3,478 replicates of control wells. Gene expression data for all datasets was collected using the L1000 assay [7].

20,131 compounds are present in both datasets. 6% percent of these compounds have MoA annotations. Only 3/20,131 compounds have replicate correlation more than 90th percentile of random distribution in both modalities.

**CDRP-bio-BBBC036-Bray-CP[35]-GE [7]:**
This is a subset of the previous dataset, containing the bioactive subset of compounds. See above (CDRP-BBBC047-Bray-CP) for dataset details.

There are 2,242 and 1,917 unique compounds for CP and GE datasets, respectively.

For CP dataset, The median number of replicates for each compound in the set is 8 and there are 3,528 replicates of control wells.
For GE dataset, The median number of replicates for each compound in the set is 2 and there are 3,478 replicates of control wells.

1,916 compounds are present in both datasets. 69% percent of these compounds have MoA annotations. 131/1,916 compounds have replicate correlation more than 90th percentile of random distribution in both modalities.

**LUAD-BBBC041-Caicedo-CP [38]- GE [11]:**

The A549 cell line was used in this genetic perturbation dataset. For CP and GE, data was collected after 96 hours of lentiviral overexpression; the plate maps are available at s3://cytodata/datasets/LUAD-BBBC043-Caicedo/metadata/LUAD-BBBC043-Caicedo/platemap/. Cell Painting was performed using the Cell Painting v1 protocol [36]. Widefield imaging was performed on an ImageXPress microscope with a 20X objective with 2x2 binning, resulting in 1080x1080 pixel images. Illumination correction, segmentation, and feature measurement were performed in CellProfiler 2.2.0, per pipelines available at https://github.com/broadinstitute/imaging-platform-pipelines/tree/master/cellpainting_a549_20x_imagexpress. L1000 was performed using the protocol described in [37]

There are 593 and 529 unique alleles for CP and GE datasets, respectively.

For CP and GE datasets, the median number of replicates for each allele in the set is 8.

525 alleles are present in both datasets. 197/525 of these alleles have replicate correlation more than 90th percentile of random distribution in both modalities.

**TA-ORF-BBBC037-Rohban-CP [3] - GE:**

The U2OS cell line was used in this genetic perturbation dataset; cells were obtained from ATCC and propagated in the William Hahn lab; they were not additionally authenticated prior to this experiment.
For CP and GE, data was collected after 72 hours of lentiviral overexpression; the platemaps are available at s3://cytodata/datasets/TA-ORF-BBBC037-Rohban/metadata/TA-ORF-BBBC037-Rohban/platemap/. Cell Painting was performed per the Cell Painting v1 protocol [36]. Widefield imaging was performed on an ImageXPress microscope with a 20X objective with 2x2 binning, resulting in 1080x1080 pixel images. Illumination correction, segmentation, and feature measurement were performed in CellProfiler 2.1, per

pipelines available as supplementary files on [3]. L1000 was performed using the protocol described in [37]

There are 323 and 327 unique alleles for CP and GE datasets, respectively.

For CP dataset, the median number of replicates for each allele in the set is 5 and there are 268 replicates of control wells.

For GE dataset, the median number of replicates for each allele in the set is 2 and there are 56 replicates of control wells.

150 alleles are present in both datasets. 36/150 of these alleles have replicate correlation more than 90th percentile of random distribution in both modalities.

**LINCS-Pilot1-CP [39] - GE [40]:**

The A549 cell line (ATCC catalog #CCL-185) was used in this chemical perturbation dataset. For CP and GE, data was collected after 24 and 48 hours of compound treatment, respectively; the platemaps are available at https://github.com/broadinstitute/lincs-cell-painting/tree/74f7a0d132e75b86aa8908689f3ce21a58a9b0f6/metadata/platemaps/2016_04_01_a549_48hr_batch1/platemap. Cell Painting was performed using the Cell Painting v2 protocol [8]; this differs from the v1 protocol only in the concentrations and Alexa dye species of the conjugated phalloidin and wheat germ agglutinin dyes (see https://github.com/carpenterlab/2016_bray_natprot/wiki#changes-in-the-official-protocol-to-create-v2-bray-et-al-2016. Widefield imaging was performed on an Opera Phenix microscope with a 20X objective with 1x1 binning, resulting in 2160x2160 pixel images. Illumination correction, segmentation, and feature measurement were performed in CellProfiler 2.2.0, per pipelines available at https://github.com/broadinstitute/imaging-platform-pipelines/tree/master/cellpainting_a549_20x_phenix_bin1. L1000 was performed using the protocol described in [7].

There are 1,570 unique compounds across 7 doses for CP dataset. There are 1,402 unique compounds across 7 doses for GE dataset. A small fraction compounds have fewer than 7 doses.

There are 9,394 and 8,369 unique compounds-dose for CP and GE datasets, respectively.

For CP dataset, the median number of replicates for each compound in the set is 5 and there are 3,264 replicates of control wells.

For GE dataset, the median number of replicates for each compound in the set is 3 and there are 1,485 replicates of control wells.

6984 compound-dose pairs are present in both datasets. 100% of these compounds have MoA annotations.

Among 6984 unique compounds-dose overlapping compounds, 1140 compound-dose have replicate correlation more than 90th percentile of random distribution in both modalities.

| Datasets | | | | | |
|---|---|---|---|---|---|
| Name | Cell Line | Perturbation Type | $N_p/N_d/N_r$ CP | $N_p/N_d/N_r$ GE | $N_p$ Intersection |
| CDRP-BBBC047-Bray | U2OS | chemical | 30,430/1/4 | 21,782/1/3 | 20,131 |
| CDRP-bio-BBBC036-Bray | U2OS | chemical | 2,242/1/8 | 1,917/1/2 | 1,916 |
| LUAD-BBBC041-Caicedo | A549 | genetic | 593/-/8 | 529/-/8 | 525 |
| TA-ORF-BBBC037-Rohban | U2OS | genetic | 323/-/5 | 327/-/2 | 150 |
| LINCS-Pilot1 | A549 | chemical | 1,570/7/ 5 | 1,402/7/3 | 6,984 |

**Supplementary Table. 1.** GE and CP dataset descriptions. $N_p$ denotes the number of perturbations. $N_d$ denotes the number of doses. $N_r$ denotes the number of replicates.

### Supplementary 2. Data Quality: Replicate reproducibility

**CDRP-BBBC047-Bray:**

CP

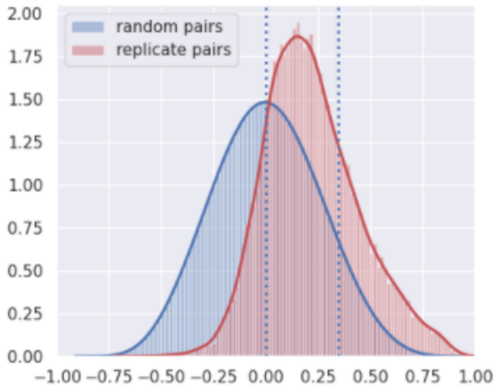


GE

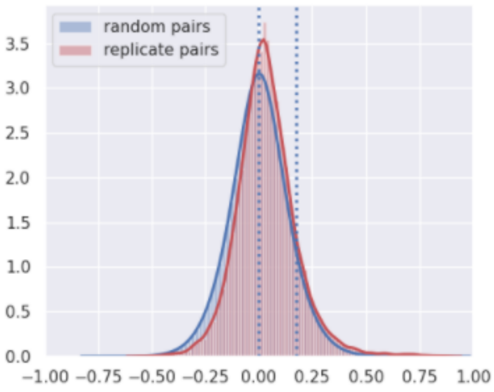


**CDRPBIO-BBBC036-Bray:**

CP

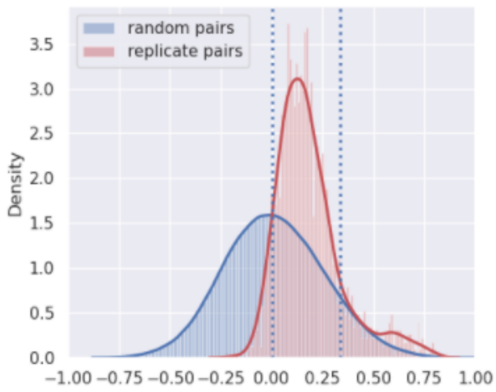


GE

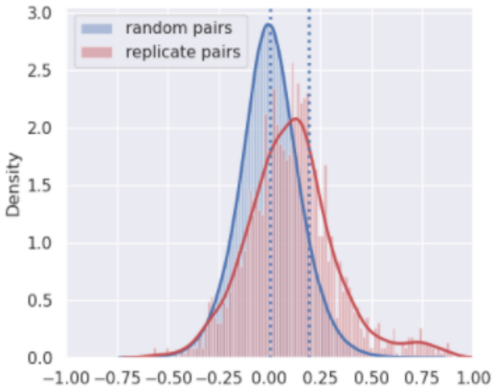


**LUAD-BBBC041-Caicedo-CP-GE :**

CP

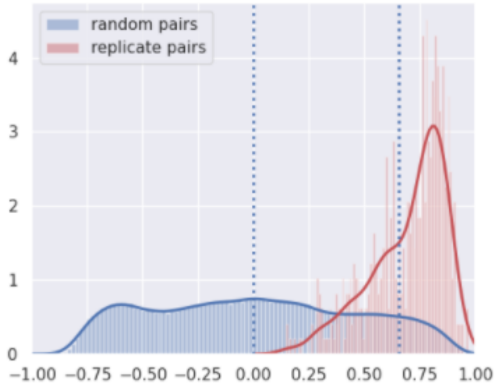


GE

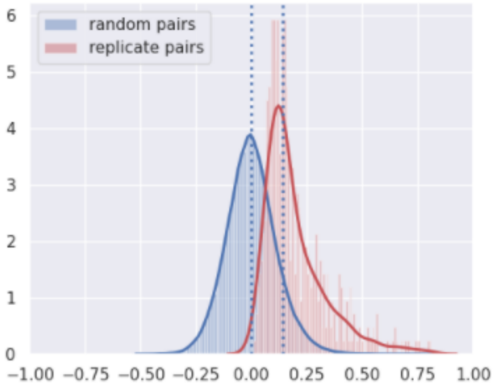

**TA-ORF-BBBC037-Rohban-CP-GE :**

| CP | GE |
|---|---|

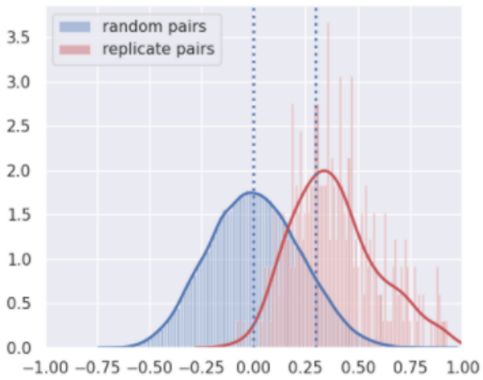


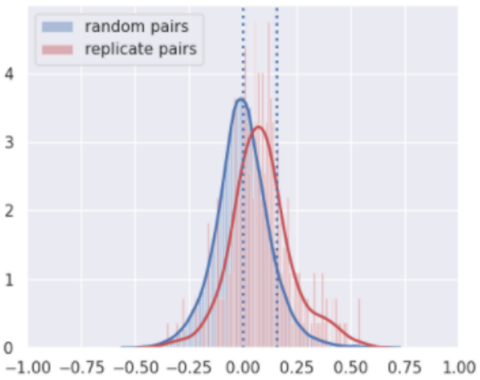


**LINCS-Pilot1-CP-GE :**

- CP - All doses together (Metadata_pert_id)
- GE - All doses together (pert_id)

| CP | GE |
|---|---|

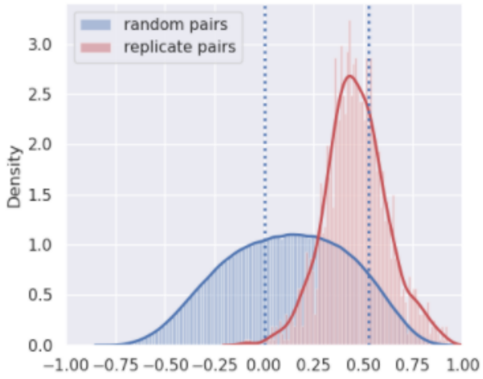


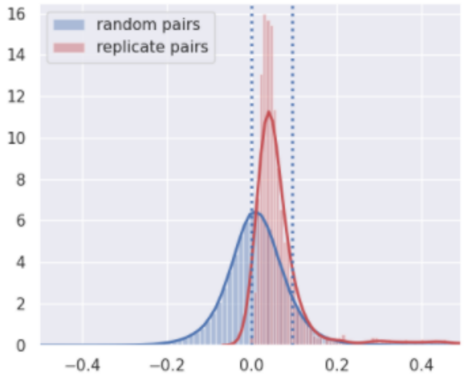


- CP - each sample/dose (Metadata_pert_id_dose)
- GE - each sample/dose (pert_id_dose)

| CP | GE |
|---|---|

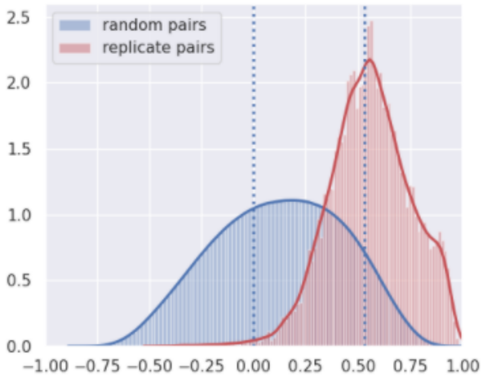


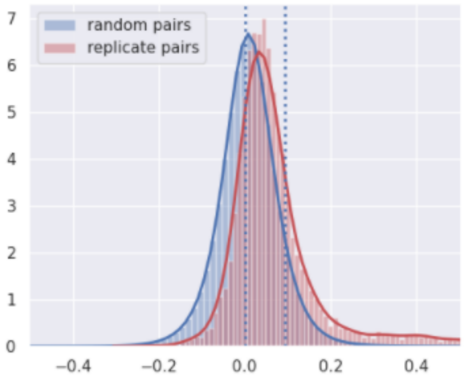

**Supplementary Figure 1.** To inspect the quality of each dataset, we calculate the consistency of profiles across different replicates of the same perturbation as follows. We standardized the profiles per plate to have zero mean and unit variance. Next, we calculated the Pearson correlation coefficient between each pair of profiles for the same perturbation (red curve) and for different perturbations (blue curve). Dotted vertical lines are shown at zero and 90th percentile of the random pairs (blue) distribution.

**Supplementary 3. Top 50 highly predictable L1000 genes by Cell Painting morphological features**

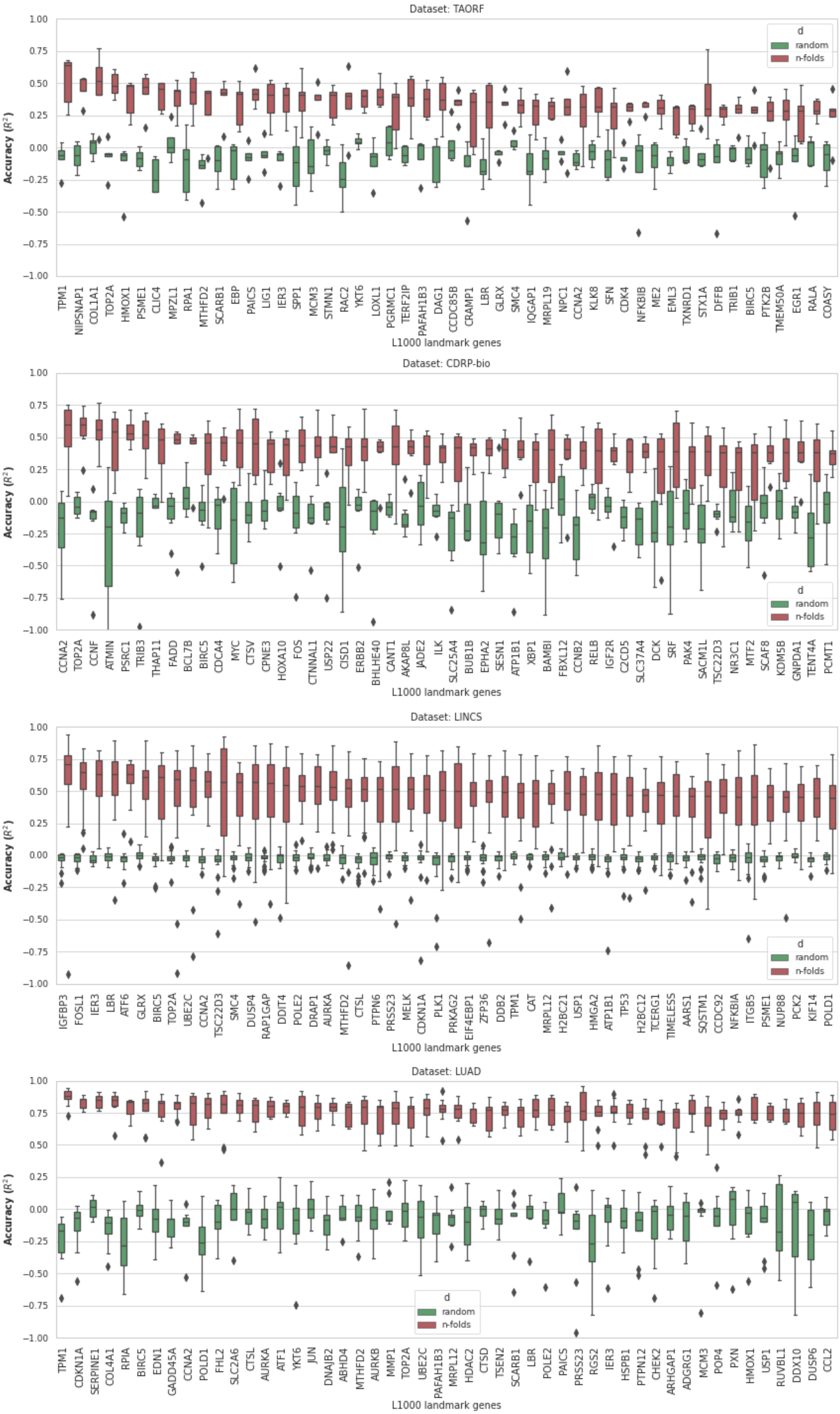
Dataset: TAORF
Accuracy ($R^2$)
1.00
0.75
0.50
0.25
0.00
−0.25
−0.50
−0.75
−1.00
d
random
n-folds
TPM1 NIPSNAP1 COL1A1 TOP2A HMOX1 PSME1 CLIC4 MPZL1 RPA1 MTHFD2 SCARB1 EBP PAICS LIG1 IER3 SPP1 MCM3 STMN1 RAC2 YKT6 LOXL1 PGRMC1 TERF2IP PAFAH1B3 DAG1 CCDC85B CRAMP1 LBR GLRX SMC4 IQGAP1 MRPL19 NPC1 CCNA2 KLK8 SFN CDK4 NFKBIB ME2 EML3 TXNRD1 STX1A DFFB TRIB1 BIRC5 PTK2B TMEM50A EGR1 RALA COASY
L1000 landmark genes
Dataset: CDRP-bio
Accuracy ($R^2$)
1.00
0.75
0.50
0.25
0.00
−0.25
−0.50
−0.75
−1.00
d
random
n-folds
CCNA2 TOP2A CCNF ATMIN PSRC1 TRIB3 THAP11 FADD BCL7B BIRC5 CDCA4 MYC CTSV CPNE3 HOXA10 FOS CTNNAL1 USP22 CISD1 ERBB2 BHLHE40 CANT1 AKAP8L JADE2 ILK SLC25A4 BUB1B EPHA2 SESN1 ATP1B1 XBP1 BAMBI FBXL12 CCNB2 RELB IGF2R C2CD5 SLC37A4 DCK SRF PAK4 SACM1L TSC22D3 NR3C1 MTF2 SCAF8 KDM5B GNPDA1 TENT4A PCMT1
L1000 landmark genes
Dataset: LINCS
Accuracy ($R^2$)
1.00
0.75
0.50
0.25
0.00
−0.25
−0.50
−0.75
−1.00
d
random
n-folds
IGFBP3 FOSL1 IER3 LBR ATF6 GLRX BIRC5 TOP2A UBE2C CCNA2 TSC22D3 SMC4 DUSP4 RAP1GAP DDIT4 POLE2 DRAP1 AURKA MTHFD2 CTSL PTPN6 PRSS23 MELK CDKN1A PLK1 PRKAG2 EIF4EBP1 ZFP36 DDB2 TPM1 CAT MRPL12 H2BC21 USP1 HMGA2 ATP1B1 TP53 H2BC12 TCERG1 TIMELESS AARS1 SQSTM1 CCDC92 NFKBIA ITGB5 PSME1 NUP88 PCK2 KIF14 POLD1
L1000 landmark genes
Dataset: LUAD
Accuracy ($R^2$)
1.00
0.75
0.50
0.25
0.00
−0.25
−0.50
−0.75
−1.00
d
random
n-folds
TPM1 CDKN1A SERPINE1 COL4A1 RPIA BIRC5 EDN1 GADD45A CCNA2 POLD1 FHL2 SLC2A6 CTSL AURKA ATF1 YKT6 JUN DNAJB2 ABHD4 MTHFD2 AURKB MMP1 TOP2A UBE2C PAFAH1B3 MRPL12 HDAC2 CTSD TSEN2 SCARB1 LBR POLE2 PAICS PRSS23 RGS2 IER3 HSPB1 PTPN12 CHEK2 ARHGAP1 ADGRG1 MCM3 POP4 PXN HMOX1 USP1 RUVBL1 DDX10 DUSP6 CCL2
L1000 landmark genes

**Supplementary Figure 2.** Prediction of L1000 mRNA levels by Cell Painting features: for each dataset, the distribution of MLP baseline prediction scores for the ordered top-50 landmark genes with the highest $R^2$ median prediction scores are shown. Each (red) boxplot represents a set of k $R^2$ values that result from running k-fold cross-validation for each corresponding landmark gene. The green boxplots are the corresponding null distributions – here we apply the exact same procedure as for the blue boxplots except that we first shuffle the output. The y-axis is trimmed at -1 for clarity. Distributions are presented as boxplots, with center line being median, box limits being upper and lower quartiles and whiskers being 1.5× interquartile range; n=k=6(CDRP-bio), 5(TAORF), 9(LUAD), 25(LINCS)

**Supplementary 4. Median Prediction scores for each landmark gene across each datasets and models**

Supplementary_D.csv

**Supplementary 5. Top 50 highly predictable Cell Painting morphological features by L1000 genes**

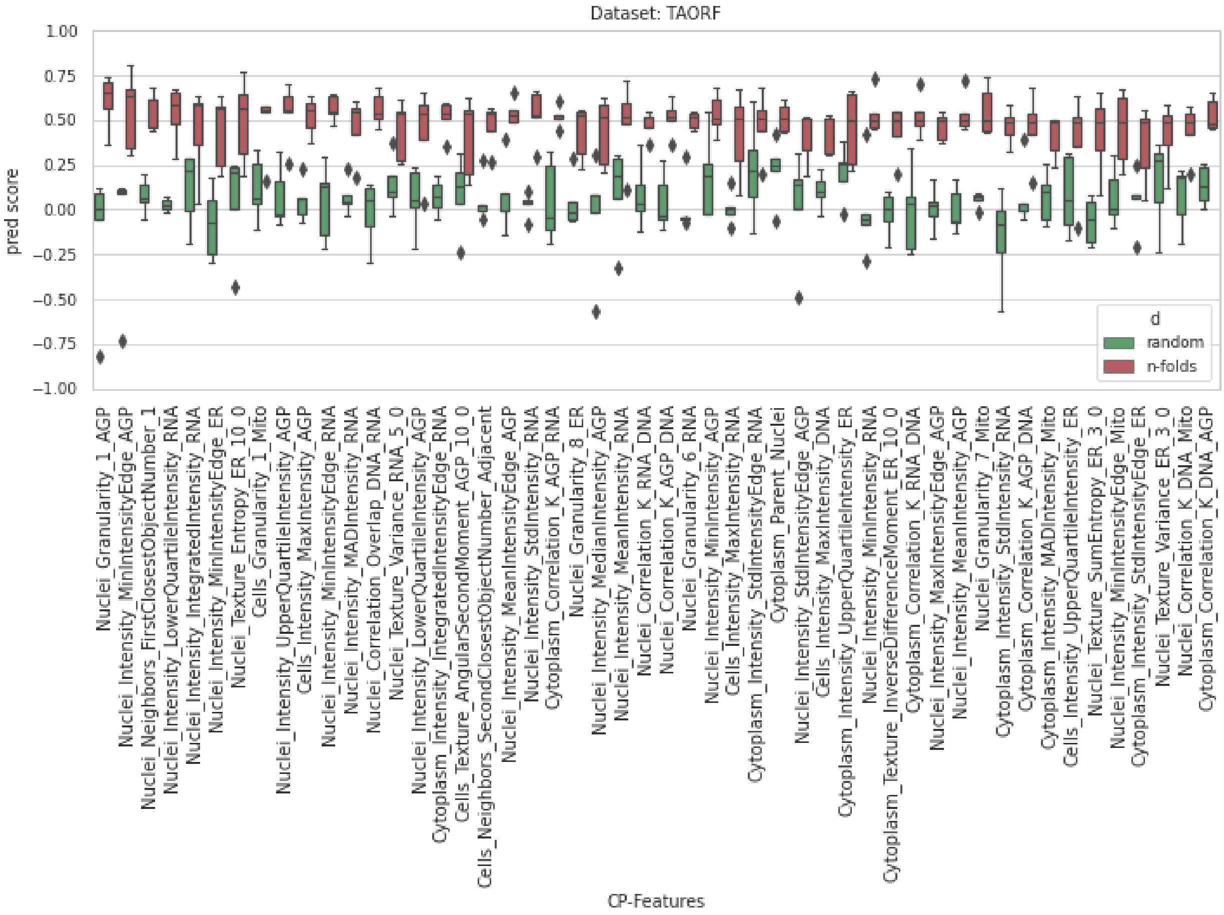
Dataset: TAORF
pred score
1.00
0.75
0.50
0.25
0.00
−0.25
−0.50
−0.75
−1.00
d
random
n-folds
Nuclei_Granularity_1_AGP
Nuclei_Intensity_MinIntensityEdge_AGP
Nuclei_Neighbors_FirstClosestObjectNumber_1
Nuclei_Intensity_LowerQuartileIntensity_RNA
Nuclei_Intensity_IntegratedIntensity_RNA
Nuclei_Intensity_MinIntensityEdge_ER
Nuclei_Texture_Entropy_ER_10_0
Cells_Granularity_1_Mito
Nuclei_Intensity_UpperQuartileIntensity_AGP
Cells_Intensity_MaxIntensity_AGP
Nuclei_Intensity_MinIntensityEdge_RNA
Nuclei_Intensity_MADIntensity_RNA
Nuclei_Correlation_Overlap_DNA_RNA
Nuclei_Texture_Variance_RNA_5_0
Nuclei_Intensity_LowerQuartileIntensity_AGP
Cytoplasm_Intensity_IntegratedIntensityEdge_RNA
Cells_Texture_AngularSecondMoment_AGP_10_0
Cells_Neighbors_SecondClosestObjectNumber_Adjacent
Nuclei_Intensity_MeanIntensityEdge_AGP
Nuclei_Intensity_StdIntensity_RNA
Cytoplasm_Correlation_K_AGP_RNA
Nuclei_Granularity_8_ER
Nuclei_Intensity_MedianIntensity_AGP
Nuclei_Intensity_MeanIntensity_RNA
Nuclei_Correlation_K_RNA_DNA
Nuclei_Correlation_K_AGP_DNA
Nuclei_Granularity_6_RNA
Nuclei_Intensity_MinIntensity_AGP
Cells_Intensity_MaxIntensity_RNA
Cytoplasm_Intensity_StdIntensityEdge_RNA
Cytoplasm_Parent_Nuclei
Nuclei_Intensity_StdIntensityEdge_AGP
Cells_Intensity_MaxIntensity_DNA
Cytoplasm_Intensity_UpperQuartileIntensity_ER
Nuclei_Intensity_MinIntensity_RNA
Cytoplasm_Texture_InverseDifferenceMoment_ER_10_0
Cytoplasm_Correlation_K_RNA_DNA
Nuclei_Intensity_MaxIntensityEdge_AGP
Nuclei_Intensity_MeanIntensity_AGP
Nuclei_Granularity_7_Mito
Cytoplasm_Intensity_StdIntensity_RNA
Cytoplasm_Correlation_K_AGP_DNA
Cytoplasm_Intensity_MADIntensity_Mito
Cells_Intensity_UpperQuartileIntensity_ER
Nuclei_Texture_SumEntropy_ER_3_0
Nuclei_Intensity_MinIntensityEdge_Mito
Cytoplasm_Intensity_StdIntensityEdge_ER
Nuclei_Texture_Variance_ER_3_0
Nuclei_Correlation_K_DNA_Mito
Cytoplasm_Correlation_K_DNA_AGP
CP-Features

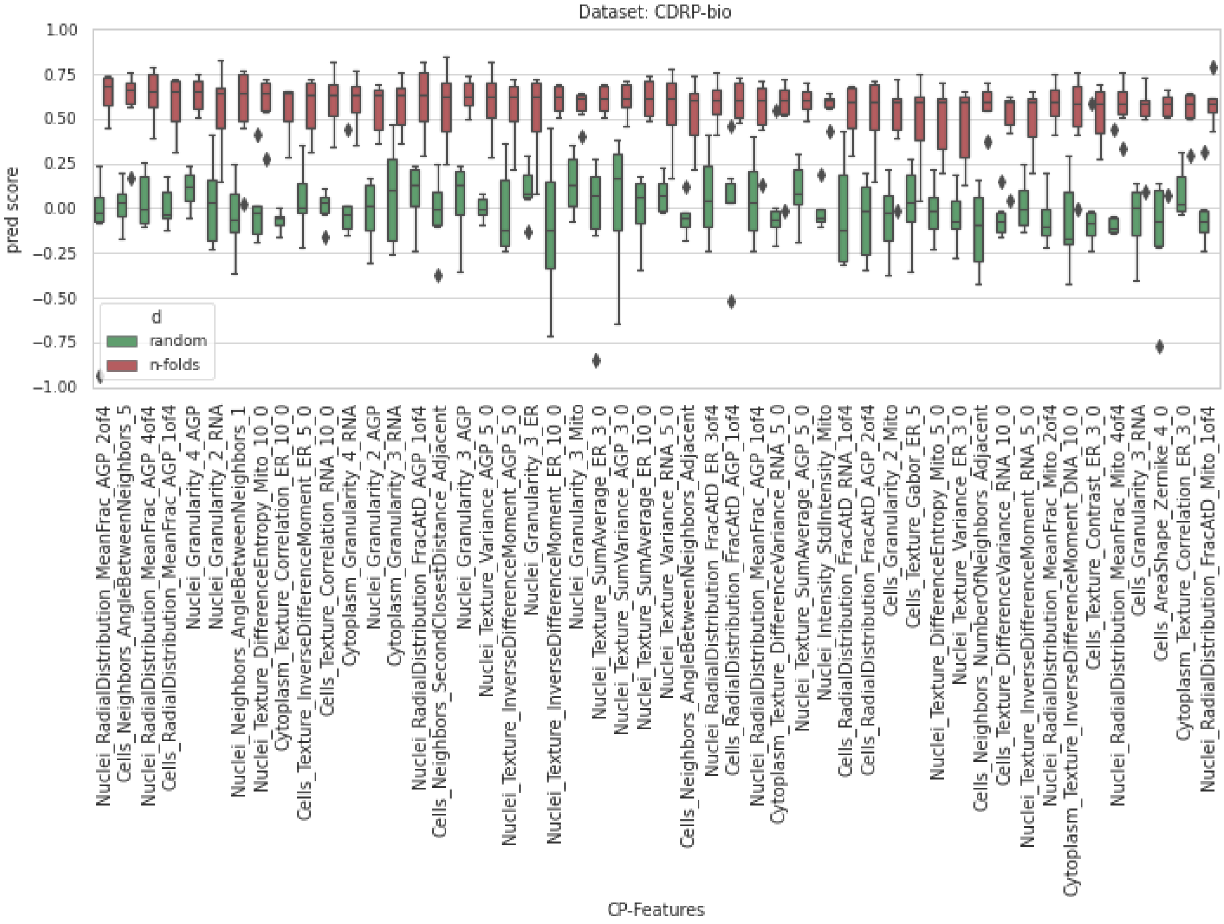
Dataset: CDRP-bio
pred score
1.00
0.75
0.50
0.25
0.00
−0.25
−0.50
−0.75
−1.00
d
random
n-folds
Nuclei_RadialDistribution_MeanFrac_AGP_2of4
Cells_Neighbors_AngleBetweenNeighbors_5
Nuclei_RadialDistribution_MeanFrac_AGP_4of4
Cells_RadialDistribution_MeanFrac_AGP_1of4
Nuclei_Granularity_4_AGP
Nuclei_Granularity_2_RNA
Nuclei_Neighbors_AngleBetweenNeighbors_1
Nuclei_Texture_DifferenceEntropy_Mito_10_0
Cytoplasm_Texture_Correlation_ER_10_0
Cells_Texture_InverseDifferenceMoment_ER_5_0
Cells_Texture_Correlation_RNA_10_0
Cytoplasm_Granularity_4_RNA
Nuclei_Granularity_2_AGP
Cytoplasm_Granularity_3_RNA
Nuclei_RadialDistribution_FracAtD_AGP_1of4
Cells_Neighbors_SecondClosestDistance_Adjacent
Nuclei_Granularity_3_AGP
Nuclei_Texture_Variance_AGP_5_0
Nuclei_Texture_InverseDifferenceMoment_AGP_5_0
Nuclei_Granularity_3_ER
Nuclei_Texture_InverseDifferenceMoment_ER_10_0
Nuclei_Granularity_3_Mito
Nuclei_Texture_SumAverage_ER_3_0
Nuclei_Texture_SumVariance_AGP_3_0
Nuclei_Texture_SumAverage_ER_10_0
Nuclei_Texture_Variance_RNA_5_0
Cells_Neighbors_AngleBetweenNeighbors_Adjacent
Nuclei_RadialDistribution_FracAtD_ER_3of4
Cells_RadialDistribution_FracAtD_AGP_1of4
Nuclei_RadialDistribution_MeanFrac_AGP_1of4
Cytoplasm_Texture_DifferenceVariance_RNA_5_0
Nuclei_Texture_SumAverage_AGP_5_0
Nuclei_Intensity_StdIntensity_Mito
Cells_RadialDistribution_FracAtD_RNA_1of4
Cells_RadialDistribution_FracAtD_AGP_2of4
Cells_Granularity_2_Mito
Cells_Texture_Gabor_ER_5
Nuclei_Texture_DifferenceEntropy_Mito_5_0
Nuclei_Texture_Variance_ER_3_0
Cells_Neighbors_NumberOfNeighbors_Adjacent
Cells_Texture_DifferenceVariance_RNA_10_0
Nuclei_Texture_InverseDifferenceMoment_RNA_5_0
Nuclei_RadialDistribution_MeanFrac_Mito_2of4
Cytoplasm_Texture_InverseDifferenceMoment_DNA_10_0
Cells_Texture_Contrast_ER_3_0
Nuclei_RadialDistribution_MeanFrac_Mito_4of4
Cells_Granularity_3_RNA
Cells_AreaShape_Zernike_4_0
Cytoplasm_Texture_Correlation_ER_3_0
Nuclei_RadialDistribution_FracAtD_Mito_1of4
CP-Features

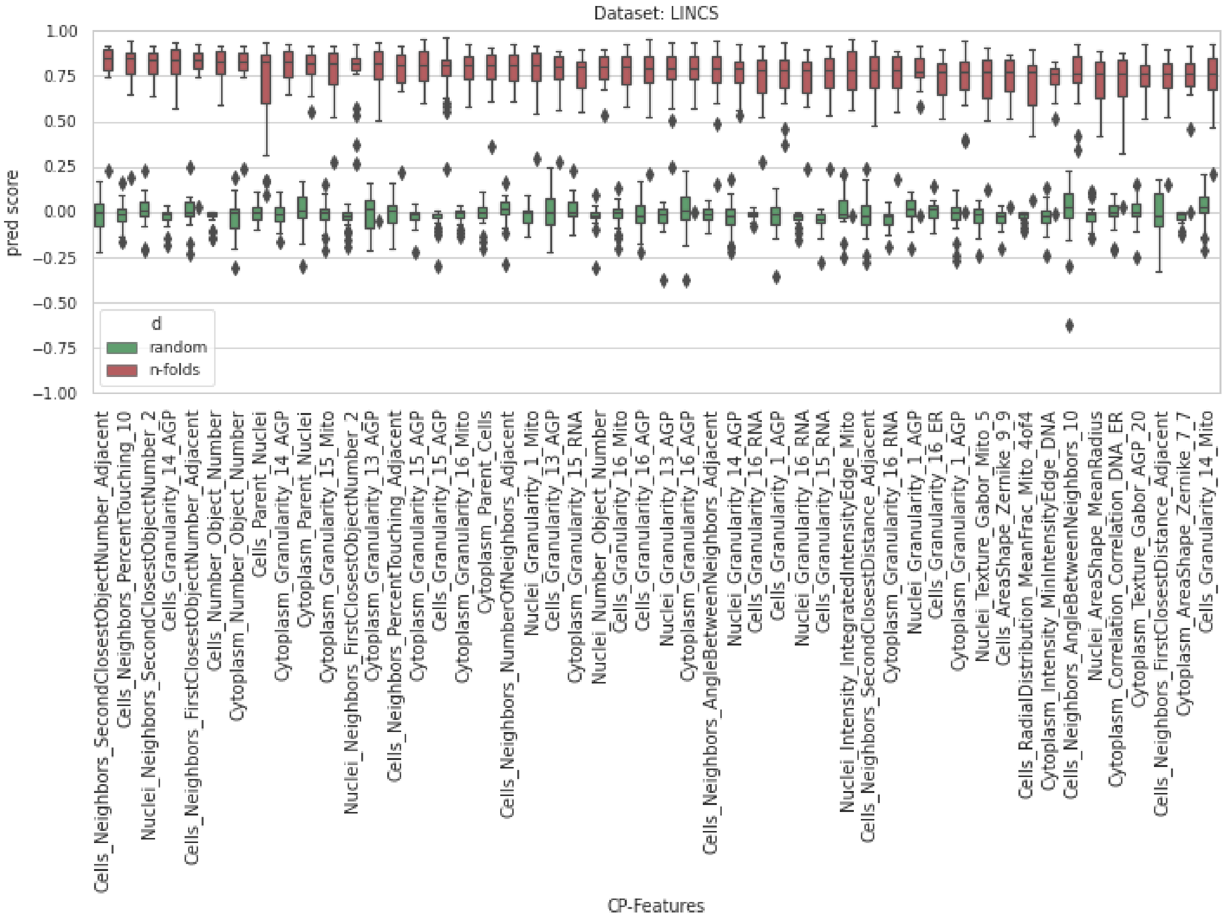
Dataset: LINCS
pred score
1.00
0.75
0.50
0.25
0.00
−0.25
−0.50
−0.75
−1.00
d
random
n-folds
Cells_Neighbors_SecondClosestObjectNumber_Adjacent
Cells_Neighbors_PercentTouching_10
Nuclei_Neighbors_SecondClosestObjectNumber_2
Cells_Granularity_14_AGP
Cells_Neighbors_FirstClosestObjectNumber_Adjacent
Cells_Number_Object_Number
Cytoplasm_Number_Object_Number
Cells_Parent_Nuclei
Cytoplasm_Granularity_14_AGP
Cytoplasm_Parent_Nuclei
Cytoplasm_Granularity_15_Mito
Nuclei_Neighbors_FirstClosestObjectNumber_2
Cytoplasm_Granularity_13_AGP
Cells_Neighbors_PercentTouching_Adjacent
Cytoplasm_Granularity_15_AGP
Cells_Granularity_15_AGP
Cytoplasm_Granularity_16_Mito
Cytoplasm_Parent_Cells
Cells_Neighbors_NumberOfNeighbors_Adjacent
Nuclei_Granularity_1_Mito
Cells_Granularity_13_AGP
Cytoplasm_Granularity_15_RNA
Nuclei_Number_Object_Number
Cells_Granularity_16_Mito
Cells_Granularity_16_AGP
Nuclei_Granularity_13_AGP
Cytoplasm_Granularity_16_AGP
Cells_Neighbors_AngleBetweenNeighbors_Adjacent
Nuclei_Granularity_14_AGP
Cells_Granularity_16_RNA
Cells_Granularity_1_AGP
Nuclei_Granularity_16_RNA
Cells_Granularity_15_RNA
Nuclei_Intensity_IntegratedIntensityEdge_Mito
Cells_Neighbors_SecondClosestDistance_Adjacent
Cytoplasm_Granularity_16_RNA
Nuclei_Granularity_1_AGP
Cells_Granularity_16_ER
Cytoplasm_Granularity_1_AGP
Nuclei_Texture_Gabor_Mito_5
Cells_AreaShape_Zernike_9_9
Cells_RadialDistribution_MeanFrac_Mito_4of4
Cytoplasm_Intensity_MinIntensityEdge_DNA
Cells_Neighbors_AngleBetweenNeighbors_10
Nuclei_AreaShape_MeanRadius
Cytoplasm_Correlation_Correlation_DNA_ER
Cytoplasm_Texture_Gabor_AGP_20
Cells_Neighbors_FirstClosestDistance_Adjacent
Cytoplasm_AreaShape_Zernike_7_7
Cells_Granularity_14_Mito
CP-Features

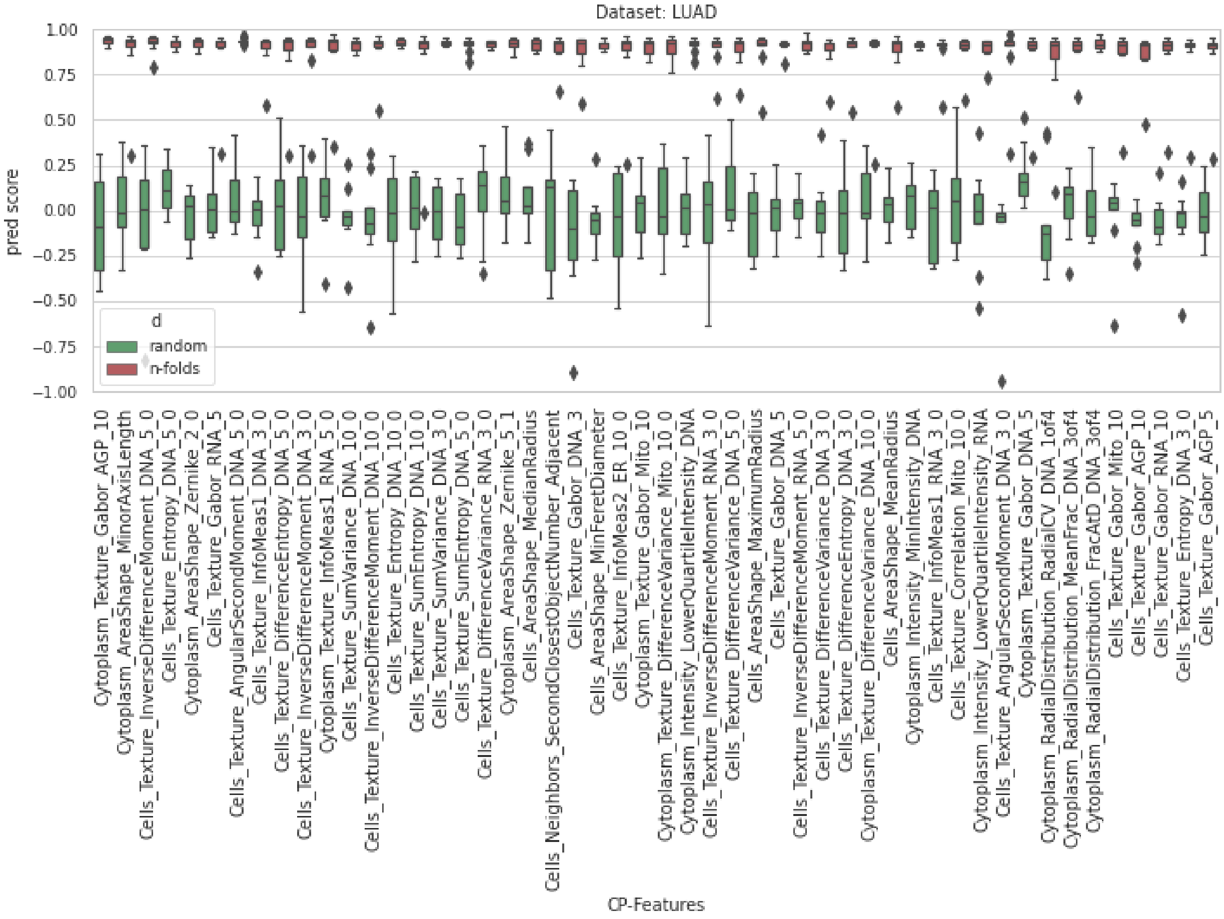
Dataset: LUAD
pred score
1.00
0.75
0.50
0.25
0.00
−0.25
−0.50
−0.75
−1.00
d
random
n-folds
Cytoplasm_Texture_Gabor_AGP_10
Cytoplasm_AreaShape_MinorAxisLength
Cells_Texture_InverseDifferenceMoment_DNA_5_0
Cells_Texture_Entropy_DNA_5_0
Cytoplasm_AreaShape_Zernike_2_0
Cells_Texture_Gabor_RNA_5
Cells_Texture_AngularSecondMoment_DNA_5_0
Cells_Texture_InfoMeas1_DNA_3_0
Cells_Texture_DifferenceEntropy_DNA_5_0
Cells_Texture_InverseDifferenceMoment_DNA_3_0
Cytoplasm_Texture_InfoMeas1_RNA_5_0
Cells_Texture_SumVariance_DNA_10_0
Cells_Texture_InverseDifferenceMoment_DNA_10_0
Cells_Texture_Entropy_DNA_10_0
Cells_Texture_SumEntropy_DNA_10_0
Cells_Texture_SumVariance_DNA_3_0
Cells_Texture_SumEntropy_DNA_5_0
Cells_Texture_DifferenceVariance_RNA_3_0
Cytoplasm_AreaShape_Zernike_5_1
Cells_AreaShape_MedianRadius
Cells_Neighbors_SecondClosestObjectNumber_Adjacent
Cells_Texture_Gabor_DNA_3
Cells_AreaShape_MinFeretDiameter
Cells_Texture_InfoMeas2_ER_10_0
Cytoplasm_Texture_Gabor_Mito_10
Cytoplasm_Texture_DifferenceVariance_Mito_10_0
Cytoplasm_Intensity_LowerQuartileIntensity_DNA
Cells_Texture_InverseDifferenceMoment_RNA_3_0
Cells_Texture_DifferenceVariance_DNA_5_0
Cells_AreaShape_MaximumRadius
Cells_Texture_Gabor_DNA_5
Cells_Texture_InverseDifferenceMoment_RNA_5_0
Cells_Texture_DifferenceVariance_DNA_3_0
Cells_Texture_DifferenceEntropy_DNA_3_0
Cytoplasm_Texture_DifferenceVariance_DNA_10_0
Cells_AreaShape_MeanRadius
Cytoplasm_Intensity_MinIntensity_DNA
Cells_Texture_InfoMeas1_RNA_3_0
Cells_Texture_Correlation_Mito_10_0
Cytoplasm_Intensity_LowerQuartileIntensity_RNA
Cells_Texture_AngularSecondMoment_DNA_3_0
Cytoplasm_Texture_Gabor_DNA_5
Cytoplasm_RadialDistribution_RadialCV_DNA_1of4
Cytoplasm_RadialDistribution_MeanFrac_DNA_3of4
Cytoplasm_RadialDistribution_FracAtD_DNA_3of4
Cells_Texture_Gabor_Mito_10
Cells_Texture_Gabor_AGP_10
Cells_Texture_Gabor_RNA_10
Cells_Texture_Entropy_DNA_3_0
Cells_Texture_Gabor_AGP_5
CP-Features

**Supplementary Figure 3.** Prediction of each Cell Painting feature by L1000 mRNA levels: for each dataset, the distribution of MLP baseline prediction scores for the ordered top-50 Cell Painting features with the highest $R^2$ median prediction scores are shown. Each (red) boxplot represents a set of k $R^2$ values that result from running k-fold cross-validation for each corresponding Cell Painting feature. The green boxplots are the corresponding null distributions – here we apply the exact same procedure as for the blue boxplots except that we first shuffle the output. The y-axis is trimmed at -1 for clarity. Distributions are presented as boxplots, with center line being median, box limits being upper and lower quartiles and whiskers being 1.5× interquartile range; n=k=6(CDRP-bio), 5(TAORF), 9(LUAD), 25(LINCS).
